# Tumor necrosis factor α modulates excitatory and inhibitory neurotransmission in a concentration-dependent manner

**DOI:** 10.1101/2022.04.14.487444

**Authors:** Dimitrios Kleidonas, Matthias Kirsch, Geoffroy Andrieux, Dietmar Pfeifer, Melanie Boerries, Andreas Vlachos

## Abstract

Microglia, the brain’s resident immune cells, have been implicated in important brain functions, such as synaptic transmission and plasticity. The pro-inflammatory cytokine tumor necrosis factor α (TNFα), which is produced and secreted by microglia, has been linked to the expression of synaptic plasticity in neurons. However, the role of TNFα-mediated activation of microglia has not been addressed in this context. Here, we assessed concentration-dependent effects of TNFα on the balance of synaptic excitation/inhibition and the activation of microglia using mouse organotypic entorhino-hippocampal tissue cultures. We found that low concentrations of TNFα enhanced excitatory synaptic strength while not affecting inhibitory neurotransmission. At higher concentrations, TNFα increased inhibitory neurotransmission without affecting excitatory synaptic strength. Both low and high concentrations of TNFα induced the synaptic accumulation of GluA1-containing AMPA receptors, suggesting that a high concentration of TNFα exerts a homeostatic effect on excitatory neurotransmission that prevents synaptic strengthening. Consistent with this, high, but not low, concentrations of TNFα activated microglia. Moreover, high concentrations of TNFα enhanced excitatory neurotransmission in microglia-depleted tissue cultures. These findings extend our knowledge on the role of TNFα on synaptic plasticity by demonstrating concentration-dependent effects on excitatory and inhibitory neurotransmission. They reveal a TNFα-mediated negative feedback mechanism on excitatory neurotransmission that is dependent on the activation of microglia, thereby emphasizing their role as gatekeepers of TNFα-mediated plasticity and homeostasis.

## INTRODUCTION

Inflammatory responses that occur in the central nervous system (CNS) after exposure to pathological stimuli, such as infections or injuries to the brain, are characterized by activation of microglia, which are the dominant immune cells localized to the brain, and increased cytokine levels [1, 2]. While this inflammatory response supports the elimination of pathogens and restricts neural damage, persistent neuroinflammatory processes can lead to severe pathological conditions, including alterations in neural plasticity, and neurodegeneration [3–6]. Numerous studies have shown that cytokines are constantly produced at low concentrations under physiological conditions in the brain [7, 8], and microglia are major sources of their production [9–12]. Several microglial pro- and anti-inflammatory cytokines have been implicated in physiological brain functions, such as synaptic transmission and plasticity [8, 13–15]. Among them, tumor necrosis factor α (TNFα) has been reported to have complex effects on the ability of neurons to express synaptic plasticity [16–26].

Long-term potentiation (LTP) induced by tetanic stimulation (1 sec, 100 Hz) was not blocked by TNFα in the rat dentate gyrus [20]. Mice lacking TNF receptors showed defects in the long-term depression, but not LTP, of Schaffer collateral CA1 synapses [17]. Recent work has also demonstrated that TNFα promotes heterosynaptic plasticity in acute hippocampal slices [27, 28]. In some studies, TNFα has been linked to changes in inhibitory neurotransmission [29, 30], however, other studies failed to demonstrate the involvement of TNFα in inhibitory synaptic plasticity [31]. Moreover, TNFα seems to mediate homeostatic plasticity of excitatory and inhibitory neurotransmission, a form of plasticity that maintains neural activity within a dynamic range through negative feedback mechanisms [23–25]. Our recent work revealed concentration-dependent effects of TNFα, whereby low concentrations promoted the ability of neurons to express synaptic plasticity and higher concentrations blocked LTP induction [22]. In this context, we proposed that alterations in LTP of excitatory neurotransmission may resemble TNFα-mediated homeostatic synaptic plasticity, which returns excitatory synaptic strength back to baseline after initial potentiation [32]. However, it remains unclear how TNFα could have mediated these concentration-dependent metaplastic effects. Specifically, the role of activated microglia in TNFα-mediated synaptic plasticity has not been previously considered.

To investigate its role in structural and functional synaptic plasticity, we tested for concentration-dependent effects of TNFα on excitatory and inhibitory neurotransmission. Accordingly, the role of TNFα-mediated microglia activation was probed in microglia-depleted organotypic tissue cultures containing the entorhinal cortex and the hippocampus (Figure 1A).

**Figure 1:**
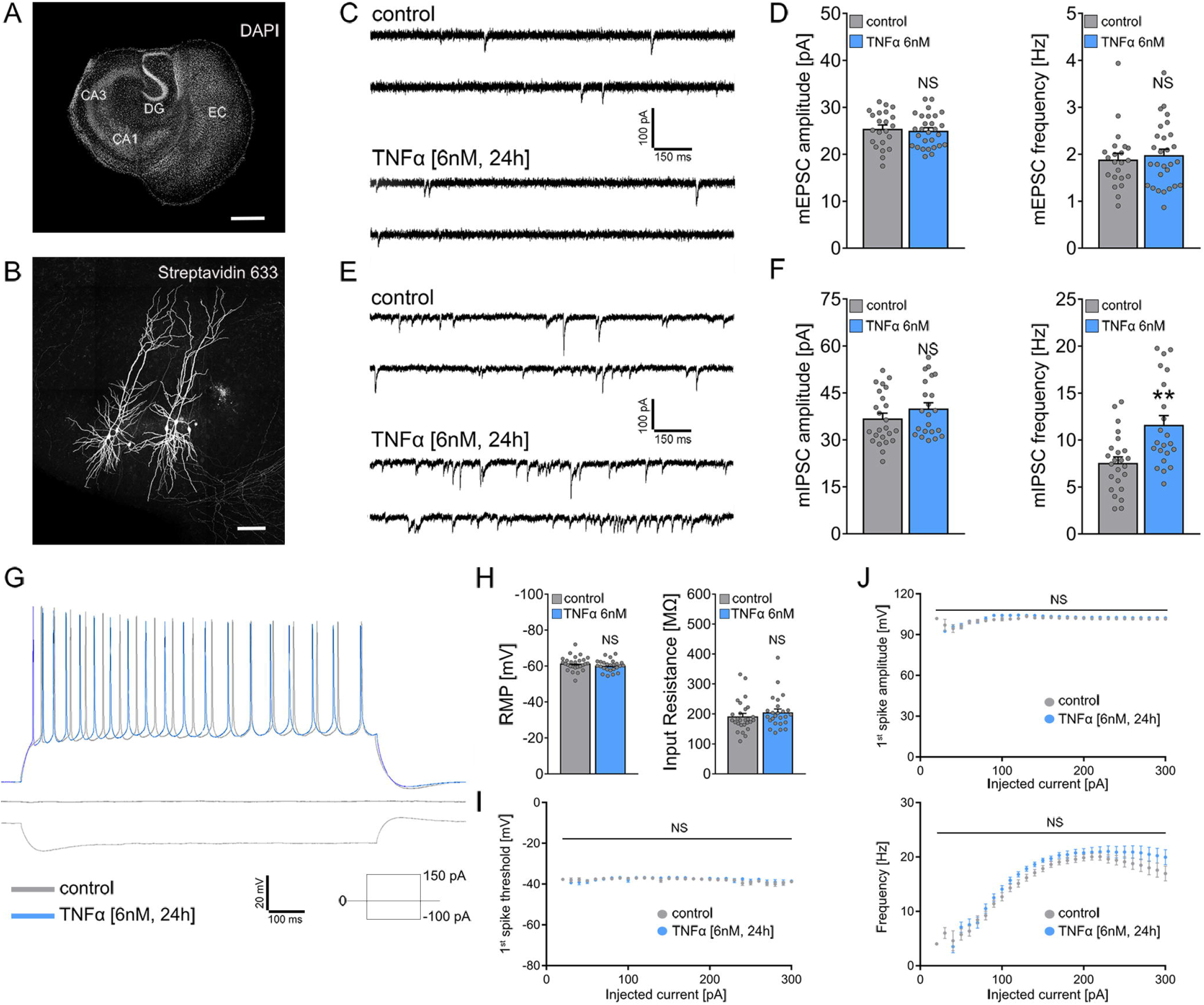
Effect of TNFα on synaptic and intrinsic cellular properties of CA1 pyramidal neurons. **(A)** Overview of a mouse organotypic entorhino-hippocampal tissue culture stained with DAPI (EC, entorhinal cortex; DG, dentate gyrus; CA1, Cornu Ammonis 1; CA3, Cornu Ammonis 3). Scale bar 500 μm. **(B)** Recorded and post-hoc identified CA1 pyramidal neurons (streptavidin 633; white). Scale bar 100 μm. **(C, D)** Sample traces and group data of AMPA receptor-mediated miniature excitatory postsynaptic currents (mEPSCs) recorded from CA1 pyramidal neurons of tissue cultures treated with vehicle-only or TNFα [6 nM, 24]. (control, n = 22 cells from 7 cultures; TNFα, n = 28 cells from 8 cultures; Mann-Whitney test). **(E, F)** Sample traces and group data of GABA receptor-mediated miniature inhibitory postsynaptic currents (mIPSCs) recorded from CA1 pyramidal neurons of tissue cultures treated with vehicle-only or TNFα [6 nM, 24]. (control, n = 24 cells from 5 cultures; TNFα, n = 22 cells from 5 cultures; Mann-Whitney test). **(G)** Sample traces of input-output properties recordings from CA1 pyramidal neurons of tissue cultures treated with vehicle-only or TNFα [6 nM, 24]. **(H)** Group data for resting membrane potentials (RMP) and input resistances in the respective groups. (control, n = 27 cells from 6 cultures; TNFα, n = 25 cells from 6 cultures; Mann-Whitney test). **(I, J)** Basic properties of action potentials recorded from CA1 pyramidal neurons of tissue cultures treated with vehicle-only or TNFα [6 nM, 24]. (control, n = 27 cells from 6 cultures; TNFα, n = 25 cells from 6 cultures; Two-way ANOVA). Values represent mean ± s.e.m., gray dots indicate individual data points (**, p<0.01; NS, not significant).

## Materials and Methods

### Ethics statement

Mice were maintained in a 12 h light/dark cycle with food and water available *ad libitum*. Every effort was made to minimize distress and pain of animals. All experimental procedures were carried out according to German animal welfare legislation and approved by the appropriate animal welfare committee and the animal welfare officer of Freiburg University.

### Animals

Wild type *C57BL/6J* and *C57BL/6-Tg(TNFa-eGFP)* [33] mice of either sex were used in this study.

### Preparation of tissue cultures

Organotypic entorhino-hippocampal tissue cultures were prepared at postnatal day 4-5 from mice of either sex as previously described [34]. The tissue cultures were transferred for cultivation onto porous (0.4 μm pore size, hydrophilic PTFE) cell culture inserts with 30 mm diameter (Millipore, Cat# PICM0RG50). The culturing medium consisted of 50 % (v/v) minimum essential medium (MEM), 25 % (v/v) basal medium eagle (BME), 25 % (v/v) heat-inactivated normal horse serum (NHS), 2 mM GlutaMAX, 0.65 % (w/v) glucose, 25 mM HEPES buffer solution, 0.1 mg/ml streptomycin, 100 U/ml penicillin and 0.15 % (w/v) bicarbonate. The pH of the culturing medium was adjusted to 7.30 and tissue cultures were incubated for at least 18 days at 35 °C in a humidified atmosphere with 5 % CO_2_. The culturing medium was replaced thrice a week.

### Pharmacology

Organotypic entorhino-hippocampal tissue cultures (≥ 18 days *in vitro*) were treated with mouse recombinant Tumor Necrosis Factor α (60 pM or 6nM; Sigma, Cat# T7539) for 24h. Only for the time course gene expression analysis TNFα remained in the culturing medium for 30 min, 3 h, 6 h, 12 h and 24 h (Figure 5D). For the depletion of microglia in tissue cultures, PLX3397 (50 nM in DMSO; Axon MedChem, Cat# 2501) was added to the culturing medium after preparation and during the whole cultivation period. The respective control tissue cultures were treated with equal volume of DMSO.

### Whole-Cell Patch-Clamp Recordings

Whole-cell patch-clamp recordings from CA1 pyramidal neurons of tissue cultures were conducted at 35°C (1 – 6 cells per culture). The basic bath solution contained 126 mM NaCl, 2.5 mM KCl, 26 mM NaHCO_3_, 1.25 mM NaH_2_PO_4_, 2 mM CaCl_2_, 2 mM MgCl_2_, 10 mM glucose and was saturated with 95 % O_2_ / 5 % CO_2_. For AMPA receptor-mediated miniature excitatory postsynaptic currents (mEPSCs) the aforementioned bath solution contained additionally 10 μM D-APV (Abcam, Cat# ab120003), 10 μM bicuculline methiodide (BMI; Tocris, Cat# 2503) and 0.5 μM tetrodotoxin (TTX; Biotrend, Cat# ARCD-0640-1). For the study of synaptic GluA1-containing AMPA receptors, mEPSCs were recorded in the presence of 10 μM 1-Naphthylacetyl spermine trihydrochloride (NASPM; Tocris, Cat# 2766). Miniature inhibitory postsynaptic current (mIPSC) recordings were conducted in bath solution containing additionally to the basic solution 10 μM D-APV, 10 μM CNQX (Abcam, Cat# ab120017) and 0.5 μM TTX. For recording of intrinsic cellular properties in current-clamp mode, the basic bath solution contained also 10 μM D-APV, 10 μM CNQX and 10 μM BMI. Patch pipettes for AMPA receptor-mediated mEPSC recordings contained 120 mM CsCH_3_SO_3_, 8 mM CsCl, 1 mM MgCl_2_, 0.4 mM EGTA, 2 mM ATP-Mg, 0.3 mM GTP-Na_2_, 10 mM PO-Creatine, 10 mM HEPES and 5mM QX-314 (pH = 7.25 with CsOH, 295 mOsm with sucrose). For miniature inhibitory postsynaptic current (mIPSC) recordings patch pipettes contained 125 mM CsCl, 5 mM NaCl, 2 mM MgCl_2_, 2 mM Mg-ATP, 0.5 mM GTP-Na_2_, 0.1 mM EGTA, and 10 mM HEPES (pH = 7.33 with CsOH, 274 mOsm with sucrose). For recording of intrinsic cellular properties patch pipettes contained 126 mM K-gluconate, 10 mM HEPES, 4 mM KCl, 4 mM ATP-Mg, 0.3 mM GTP-Na_2_, 10 mM PO-Creatine, and 0.1% (w/v) biocytin (pH 7.25 with KOH, 290 mOsm with sucrose). Miniature excitatory postsynaptic currents (mEPSCs) and miniature inhibitory postsynaptic currents (mIPSCs) were recorded at a holding potential of - 60 mV. Series resistance was monitored before and after each recording and recordings were discarded if the series resistance reached ≥30 MΩ. In current-clamp mode, IV-curves were generated by injecting 1 s square pulse currents starting at −100 pA and increasing with 10 pA steps until +300 pA current injection was reached (sweep duration: 2 s).

### Post-hoc identification of recorded neurons

After recording the intrinsic cellular properties, tissue cultures were fixed in a solution of 4 % (w/v) paraformaldehyde (PFA) and 4 % (w/v) sucrose in 0.01 M PBS for 1 h at room temperature. After fixation the tissue cultures were washed with 0.01 PBS. Then, the fixed tissue cultures were incubated for 1 h at room temperature in blocking solution consisting of 10 % (v/v) normal goat serum (NGS; Fisher Scientific, Cat# NC9270494) and 0.5 % (v/v) Triton X-100 in 0.01 M PBS. Biocytin- (Sigma-Aldrich, Cat# B4261) filled cells were stained with Alexa-633 conjugated Streptavidin (1:1000 in 0.01 M PBS with 10 % NGS and 0.1 % Triton X-100; Thermo Fisher Scientific, Cat# S21375) overnight at 4 °C (while shaking). DAPI (Thermo Fisher Scientific, Cat# 62248) staining was used to visualize cytoarchitecture (1:2000; in 0.01 M PBS for 15 min). Slices were washed three times with 0.01 M PBS, transferred and mounted onto glass slides with anti-fading mounting medium (DAKO; Agilent, Cat# S3023) for visualization. Streptavidin-stained CA1 pyramidal neurons were visualized with a Leica TCS SP8 laser scanning microscope with 20x (NA 0.75; Leica), 40x (NA 1.30; Leica) and 63x (NA 1.40; Leica) oil-submersion objectives.

### Immunohistochemistry and imaging

Tissue cultures were fixed in a solution of 4 % (w/v) paraformaldehyde (PFA) and 4 % (w/v) sucrose in 0.01 M phosphate-buffered saline (PBS) for 1 h at room temperature, followed by 2 % (w/v) PFA and 30 % (w/v) sucrose in 0.01 M PBS overnight at 4 °C for cryoprotection. 30 μm cryo-sections were prepared using a cryostat (Leica CM3050 S) and were incubated for 1 h at room temperature in blocking solution consisting of 10 % (v/v) NGS and 0.5 % (v/v) Triton X-100 in 0.01 M PBS to reduce non-specific antibody binding. After blocking, the cryo-sections were incubated for 48 h at 4 °C (while shaking) with a solution containing mouse primary antibody against GluA1 (1:1000; Synaptic Systems, Cat# 182011) in 0.01 M PBS with 10 % NGS and 0.1 % Triton X-100.

Sections were washed twice in 0.01 M PBS and incubated overnight at 4 °C with appropriate Alexa-labelled secondary antibody (goat anti-mouse Alexa plus 555-labelled secondary antibody; Invitrogen; 1:1000, Cat# A32727) in PBS with 10 % NGS, 0.1 % Triton X-100. DAPI nuclear staining was used to visualize cytoarchitecture (1:5000; in 0.01 M PBS for 15 min). Sections were washed three times with 0.01 M PBS, transferred onto glass slides and mounted for visualization with anti-fading mounting medium.

For microglia visualization a similar procedure was followed. Again, tissue cultures were fixed in a solution of 4 % (w/v) paraformaldehyde (PFA) and 4 % (w/v) sucrose in 0.01 M phosphate-buffered saline (PBS) for 1 h at room temperature but were then directly washed with and stored in 0.01 M PBS until further experimental assessment. Afterwards, the fixed tissue cultures were incubated for 1 h at room temperature in a blocking solution consisting of 10% (v/v) NGS and 0.5% (v/v) Triton X-100 in 0.01 M PBS. After blocking, the fixed tissue cultures were incubated for 48 h at 4 °C (while shaking) with a solution containing rabbit primary antibody against Iba1 (1:1000; Wako, Cat# 019-19741) in 0.01 M PBS with 10% NGS and 0.1% Triton X-100. The fixed cultures were washed twice in 0.01 M PBS and incubated overnight at 4 °C with donkey anti-rabbit Alexa488-labeled secondary antibody (1:1000; Invitrogen, Cat# A21206) in 0.01 M PBS with 10% NGS, 0.1% Triton X-100. DAPI nuclear staining was used to visualize cytoarchitecture (1:2000; in 0.01 M PBS for 15 min). Finally, the samples were washed thrice with 0.01 M PBS, transferred onto glass slides and mounted for visualization with anti-fading mounting medium. Confocal images were acquired using a Leica TCS SP8 laser scanning microscope with 20x (NA 0.75; Leica), 40x (NA 1.30; Leica) and 63x (NA 1.40; Leica) oil-submersion objectives.

### Surface biotinylation

Tissue cultures (4 per sample) were washed in cold solution, containing 126 mM NaCl, 2.5 mM KCl, 26 mM NaHCO_3_, 1.25 mM NaH_2_PO_4_, 2 mM CaCl_2_, 2 mM MgCl_2_ and 10 mM glucose. Biotin 3-sulfo-N-hydroxysuccinimide ester (1 mg/mL; Sigma, Cat# B5161) was added for 30 minutes, while shaking at 4 °C. 10 mM NH_4_Cl was added for quenching and after washing with cold ACSF, the tissue cultures were harvested in lysis solution. Tissue cultures were homogenized using a Hamilton syringe and after centrifugation the supernatant was collected. The supernatant was added to streptavidin coupled magnetic beads (Steinbrenner Laborsysteme, Cat# MD21001) and incubated overnight at 4 °C while shaking (~ 700 - 800 rpm). The beads were washed 5 times with a solution containing 1 mM EDTA, 150 mM NaCl, 50mM Tris-HCl (pH 7.5) and 1% Triton X-100. For the last washing step a solution containing 1 mM EDTA in 10 mM Tris-HCl (pH 7.5) was used. Finally, the beads were heated in sample buffer at 95 °C.

### Western blot analysis

15 μL of lysate were loaded in each well of polyacrylamide gels (ThermoFisher, Cat# XP04205BOX) and electrophoresis was performed. The samples were then transferred on PVDF membrane. For blocking, the PVDF membrane was incubated for 1h in 5 % milk TBS-T (TBS/0.1 % Tween 20 + 5 % Milk Powder). The membrane was then incubated overnight in 5 % milk TBS-T containing mouse anti-GluA1 (1:1000: Synaptic Systems, Cat# 182011) and mouse anti-GAPDH (1:2000; Novus, Cat# NB600-502) antibodies. Goat anti-HRP antibody (1:10000; Jackson ImmunoResearch; Cat# 115-035-003) was used for binding to the primary antibodies. Clarity Max ECL substrate (Biorad, Cat# 1705062) was used for detection.

### Propidium iodide stainings

Tissue cultures were incubated with propidium iodide (1 μg/mL; Invitrogen, Cat# P3566) for 1h, washed in PBS (0.1 M, pH 7.4) and fixed as described above. Tissue cultures treated with NMDA (50 μM; Tocris, Cat# 0114) for 24 h served as positive controls. Cell nuclei were stained with DAPI and the cultures were mounted as described earlier.

### Transcriptome Analysis

Tissue cultures (n=3) were transferred as one sample into 250 μl RNAlater (ThermoFisher, Cat# AM7020) and stored at −20 °C. RNA was isolated after homogenization with TRIzol (ThermoFisher, Cat# 15596018) using the Direct-zol RNA Microprep-Kit (Zymo Research, #R2061) according to the manufacturer’s instructions. RNA was eluted in 50 μl water and precipitated with 0.75 M ammonium-acetate and 10 μg glycogen (ThermoFisher, Cat# R0551) by adding 125 μl ethanol (100 %). Samples were incubated at −80 °C overnight and consecutively centrifuged for 30 minutes at 4 °C. Pellets were washed with 70 % ethanol, centrifuged again and dried. Finally, pellets were dissolved in water for further processing. RNA concentration and integrity was consecutively analyzed by capillary electrophoresis using a Fragment Analyser (Advanced Analytical Technologies, Inc., USA) and the Agilent RNA 6000 Pico Kit (Agilent, Cat# 5067-1513). RNA samples with RNA integrity numbers (RIN) > 8.0 were further processed with the Affymetrix WT Plus kit and hybridized to Clariom S mouse arrays (ThermoFisher, Cat# 902931) as described by the manufacturer. Briefly, labeled fragments were hybridized to arrays for 16 h at 45 °C, 60 rpm in a GeneChip™ Hybridization Oven (ThermoFisher). After washing and staining, the arrays were scanned with the Affymetrix GeneChip Scanner 3000 7G (ThermoFisher). CEL files were produced from the raw data with Affymetrix GeneChip Command Console Software Version 4.1.2 (ThermoFisher). CEL files were processed with the Oligo R package and RNA and intensity were normalized using Robust Multichip Average method. A linear-model based analysis, limma R package [35], was used to identify differentially regulated genes. An adjusted p value (Benjamini & Hochberg) below 0.05 was considered as significant. Gene-set enrichment analysis was done using GSEA 4.1.0 [36, 37], Cytoscape [38] and Custom R-scripts to produce heatmaps.

### Quantitative reverse transcription PCR (RT-qPCR)

500 ng of purified RNA (see above) was reverse transcribed (RevertAid RT Kit; ThermoFisher, Cat# K1691). The resulting cDNA was diluted 1:50 in water and subjected to RT-qPCR using the CFX384 Touch Real-Time PCR Detection System (BIO-RAD) and the ORA-qPCR Green ROX L 2X Mix (HighQu, Cat# QPD0101). Primers (see table below) were used at 200 nM each. The RT-qPCR protocol was performed as follows: 95 °C for 3 min, 40 cycles of 95 °C / 30 sec, 61.5 °C / 60 sec, melting curve: 65 °C to 95 °C, 0.5 °C increments. Analysis was conducted using the Bio-Rad CFX Maestro software package and normalization was performed with 3 house-keeping genes (*Gapdh, Alas1, Rps12*).

### Primer sequences

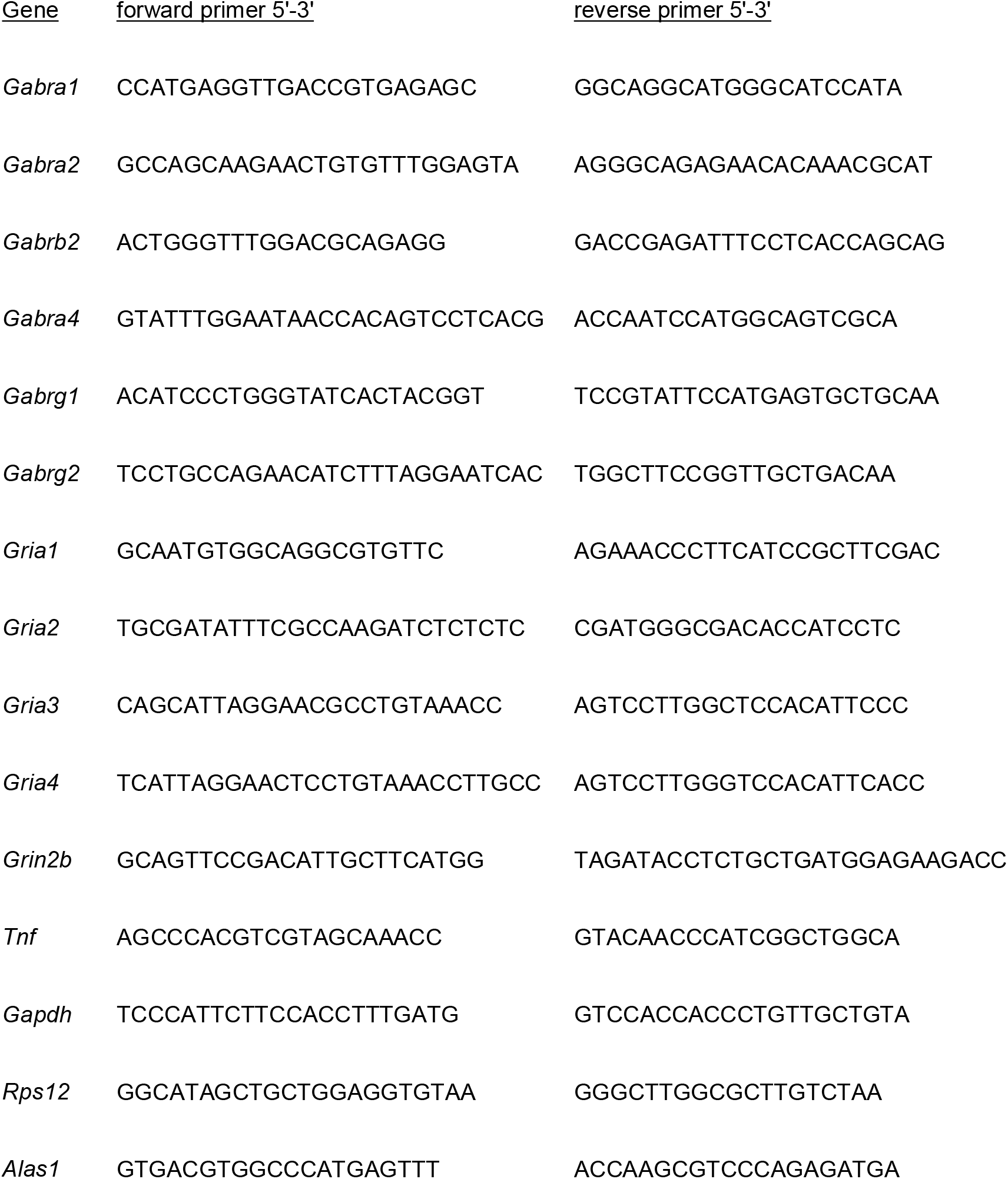

### Live-cell imaging

Live-cell imaging of heterozygous *C57BL/6-Tg(TNFa-eGFP)* cultures, either vehicle-treated or TNFα-treated, was performed at a Zeiss LSM800 microscope equipped with a 10x water-immersion objective (NA 0.3; Carl Zeiss). Filter membranes with 3 cultures were placed in a 35 mm Petri Dish containing pre-oxygenated imaging solution consisting of 50 % (v/v) MEM, 25 % (v/v) basal medium eagle, 50 mM HEPES buffer solution (25 % v/v), 0.65 % (w/v) glucose, 0.15 % (w/v) bicarbonate, 0.1 mg/ml streptomycin, 100 U/ml penicillin, 2 mM GlutaMAX, and 0.1 mM Trolox (6-Hydroxy-2,5,7,8-tetramethylchromane-2-carboxylic acid; Sigma, Cat# 238813). The cultures were kept at 35 °C during the imaging procedure. Laser intensity and detector gain were initially set to keep the fluorescent signal in a dynamic range throughout the experiment and were kept constant in all experiments. Confocal image stacks were stored as czi files.

### ELISA assay for cytokine detection

Incubation medium was collected from mature (>18 DIV) entorhino-hippocampal tissue cultures (3 per insert) treated either with vehicle-only or TNFα (60 pM or 6 nM, 24h) and stored immediately at −80 °C. In order to determine the concentration of pro-inflammatory and anti-inflammatory cytokines in the culturing medium, a V-Plex Proinflammatory Panel 1 (mouse) Kit Plus (Mesoscale Discovery #K15048G) was used. The collected incubation medium was diluted 1:1 in diluent provided with the kit. Protein detection was performed according to manufacturer’s instructions. A pre-coated plate with captured antibodies on defined spots was incubated with the diluted samples for 2 h. After washing, samples were incubated for 2 h with a solution containing electrochemiluminescent MSD SULFO-TAG detection antibodies (Mesoscale Discovery; Antibodies: anti-ms IL1β antibody #D22QP, anti-ms CXCL1 antibody #D22QT, anti-ms IL6 antibody #D22QX, anti-ms IL10 antibody, #D22QU). After washing, samples were measured with a MESO QuickPlex SQ 120 instrument (Mesoscale Discovery). The respective protein concentrations were determined using the MSD DISCOVERY WORKBENCH software (Mesoscale Discovery).

### Experimental Design and statistical analysis

Electrophysiological recordings were analyzed using Clampfit 11 of the pClamp11 software package (Molecular Devices). mEPSC and mIPSC properties were analyzed using the automated template search tool for event detection. Action potential properties were analyzed using Igor Pro 7 (Wavemetrics). Numbers and sizes of immunolabeled GluA1 clusters were assessed using the FIJI ImageJ software package (available from https://imagej.net/ImageJ) as previously described [39]. Spines of second- and third-order dendritic branches from streptavidin-labelled CA1 pyramidal neurons were analyzed after maximum intensity projection. Blinded analyses of spine density was performed manually using the FIJI ImageJ software package. For PI stainings, fluorescence intensity was analyzed in the whole tissue culture using the particle analysis of FIJI ImageJ software package. Again, the analysis was performed in a blinded manner. For the transcriptome analysis CEL-files were analyzed using the Transcriptome Analysis Software package (TAC, ThermoFisher) for differentially expressed genes and custom R-scripts to produce heatmaps. Analysis of qPCR data was performed using the Bio-Rad CFX Maestro software package and normalization was done with 3 house-keeping genes (*Gapdh, Alas1, Rps12*). Confocal image stacks of heterozygous *C57BL/6-Tg(TNFa-eGFP*) cultures were processed and analyzed using the Fiji image processing package. From each stack maximum intensity projection of 3 images was performed. Tissue culture area and the molecular layer of the dentate gyrus were manually defined as regions of interest (ROI) and the mean fluorescence intensity of each ROI was measured by an investigator blind to the experimental conditions. The protein concentrations of cytokines in the culture medium were determined using the MSD DISCOVERY WORKBENCH software (Mesoscale Discovery). Statistical comparisons were carried out using GraphPad Prism 7 (GraphPad software). For comparison of two groups Mann-Whitney test was used. In order to statistically compare three groups Kruskal-Wallis test followed by Dunn’s post hoc test was selected. For statistical evaluation of XY-plots, a two-way ANOVA test was performed.

### Digital illustrations

Figures were prepared using the Affinity Designer (Serif Europe) and the Adobe Photoshop (Adobe) graphics software. Image brightness and contrast were adjusted. Figures 9A and 10 were prepared with Biorender (www.biorender.com).

## RESULTS

### TNFα affects inhibitory but not excitatory neurotransmission

Tissue cultures (≥18 days *in vitro*; Figure 1A) were exposed to 6 nM TNFα for 24 hours (c.f.,[24, 40–42]), with vehicle-only tissue cultures used as a control. AMPA receptor-mediated miniature excitatory postsynaptic currents (mEPSCs) and GABA receptor-mediated miniature inhibitory postsynaptic currents (mIPSCs) were recorded from individual CA1 pyramidal neurons (Figure 1B). Agreeing with our previous studies, which revealed no major effects of TNFα on excitatory neurotransmission under baseline conditions [22, 25], mean mEPSC amplitude and frequency were not significantly different between vehicle-only and TNFα-treated tissue cultures (Figure 1C, D). When mIPSCs were recorded in a different set of tissue cultures, a significant increase in mean mIPSC frequency was observed in the TNFα group, whereas mIPSC amplitudes were indistinguishable from the vehicle-only control (Figure 1E, F).

We also recorded input-output properties of CA1 pyramidal neurons (Figure 1G) and found no significant differences between TNFα exposed and vehicle-only controls. Resting membrane potential, input resistance (Figure 1H) and basic properties of action potentials (Figure 1I, J) were not significantly different between controls and TNFα-treated tissue cultures. These results suggest that TNFα exposure (6 nM, 24 hours) affected inhibition while leaving excitatory neurotransmission and basic input-output properties of CA1 pyramidal neurons intact.

### TNFα promotes synaptic insertion of GluA1-containing AMPA receptors

Increased inhibition is expected to hamper the ability of neurons to express synaptic plasticity [43, 44]. In previous studies, we questioned how 6 nM TNFα could assert its plasticity-promoting effects in neural networks [22], speculating that TNFα might change AMPA receptor subunit composition without affecting baseline synaptic strength. Specifically, TNFα may increase the synaptic content of Ca^2+^-permeable, GluA2-lacking AMPA receptors, i.e., GluA1-containing AMPA receptors [41], which promote the ability of neurons to express plasticity [45].

To test this hypothesis, AMPA receptor-mediated mEPSCs were recorded from CA1 pyramidal neurons in vehicle-only and TNFα-treated (6 nM, 24 hours) tissue cultures (Figure 2A, B). Experiments were performed in the presence of 1-naphthylacetyl spermine (NASPM; 10 μM), which blocks Ca^2+^-permeable, thus, mainly GluA1-only-containing AMPA receptors. In the presence of NASPM, mEPSC frequencies were significantly reduced after TNFα treatment (Figure 2B), suggesting that a higher number of synaptic GluA1-containing AMPA receptors were found in the TNFα group.

**Figure 2:**
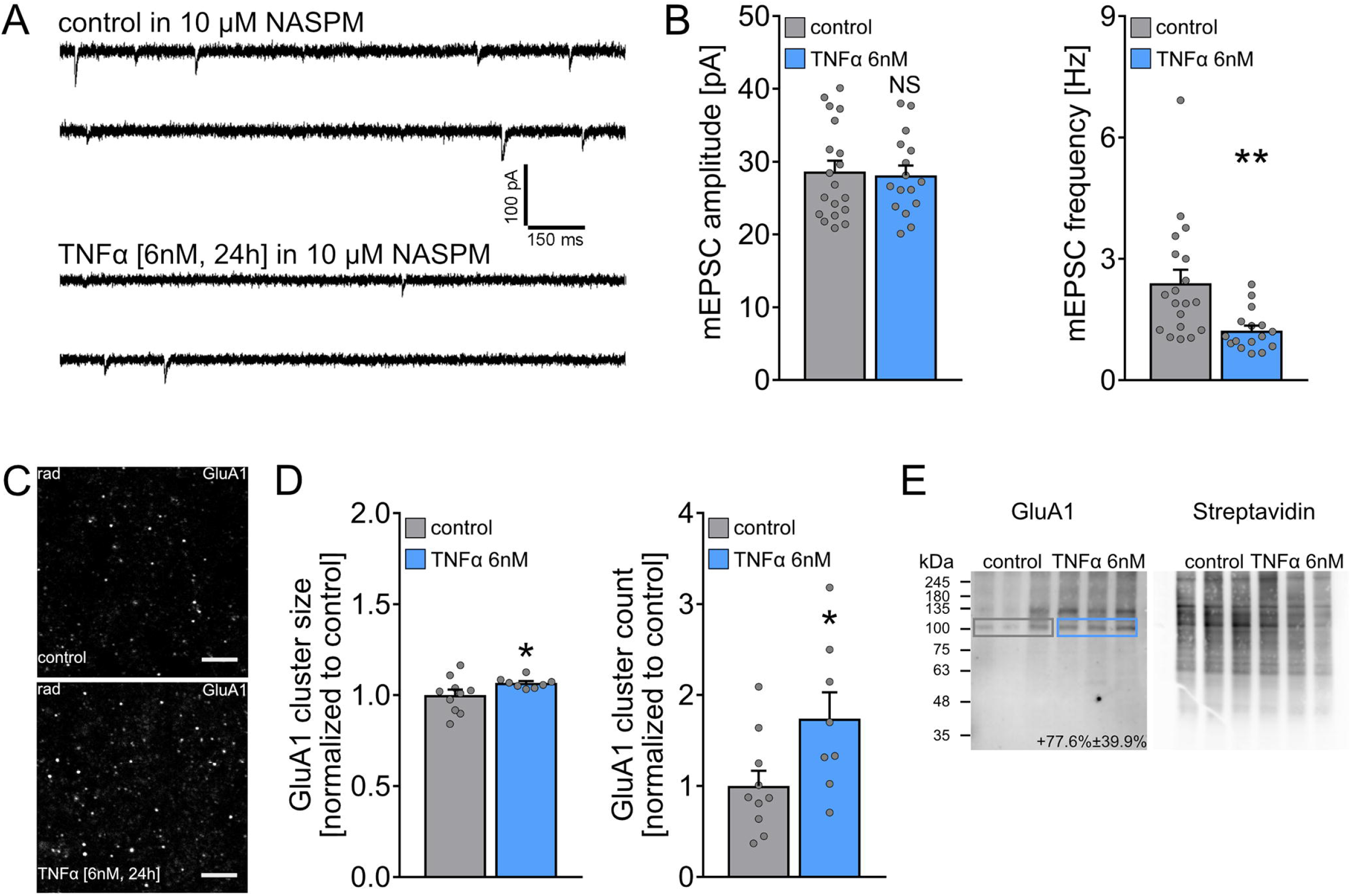
TNFα leads to synaptic accumulation of 1-Naphthylacetyl spermine sensitive (Ca^2+^-permeable) AMPA receptors. **(A, B)** Sample traces and group data of AMPA receptor-mediated miniature excitatory postsynaptic currents (mEPSCs) recorded from CA1 pyramidal neurons of tissue cultures treated with vehicle-only or TNFα [6 nM, 24] in the presence of 1-Naphthylacetyl spermine [NASPM;10 μM]. (control, n = 19 cells from 4 cultures; TNFα, n = 16 cells from 4 cultures; Mann-Whitney test). Values represent mean ± s.e.m., gray dots indicate individual data points (**, p<0.01; NS, not significant). **(C, D)**, Representative images and group data of GluA1 clusters in the stratum radiatum of the CA1 region from tissue cultures treated with vehicle-only (upper) or TNFα [6 nM, 24] (lower). (control, n = 10 cultures; TNFα, n = 8 cultures; Mann-Whitney test). Values represent mean ± s.e.m., gray dots indicate individual data points (* p<0.05). Scale bar 4 μm. **(E, left)** Western-blot detection of GluA1 after surface biotinylation of cultures treated with vehicle-only (grey rectangle) or TNFα [6 nM, 24] (blue rectangle) **(E, right)** Blot with streptavidin-HRP to detect all biotinylated proteins in lysates from cultures treated with vehicle-only or TNFα [6 nM, 24]. (control, n = 3 samples; TNFα, n = 3 samples).

To confirm these results, a different set of tissue cultures was immunostained for GluA1, and cluster sizes and numbers were assessed as described previously [39]. A significant increase in GluA1 cluster sizes and numbers was observed in the CA1 stratum radiatum in the TNFα-treated group compared to vehicle-only controls (Figure 2C, D). Consistent with these data and our electrophysiological recordings, surface biotinylation confirmed increased surface levels of GluA1 in TNFα-treated tissue cultures (Figure 2E). Given these results, we concluded that TNFα increased surface GluA1 levels and promoted the accumulation of GluA1-containing AMPA receptors at excitatory postsynapses without affecting baseline synaptic transmission.

### TNFα does not affect the number of dendritic spines of CA1 pyramidal neurons

Given the increased numbers of GluA1 clusters (cf. Figure 2C, D), we next tested whether TNFα could increase the density of dendritic spines (Figure 3). Identified side branches of main apical dendrites in CA1 stratum radiatum of recorded, biocytin-filled and post hoc-stained pyramidal neurons were analyzed in these experiments. As shown in Figure 3, we did not find any significant differences in spine numbers between vehicle-treated and TNFα-treated (6 mM, 24 hours) tissue cultures.

**Figure 3:**
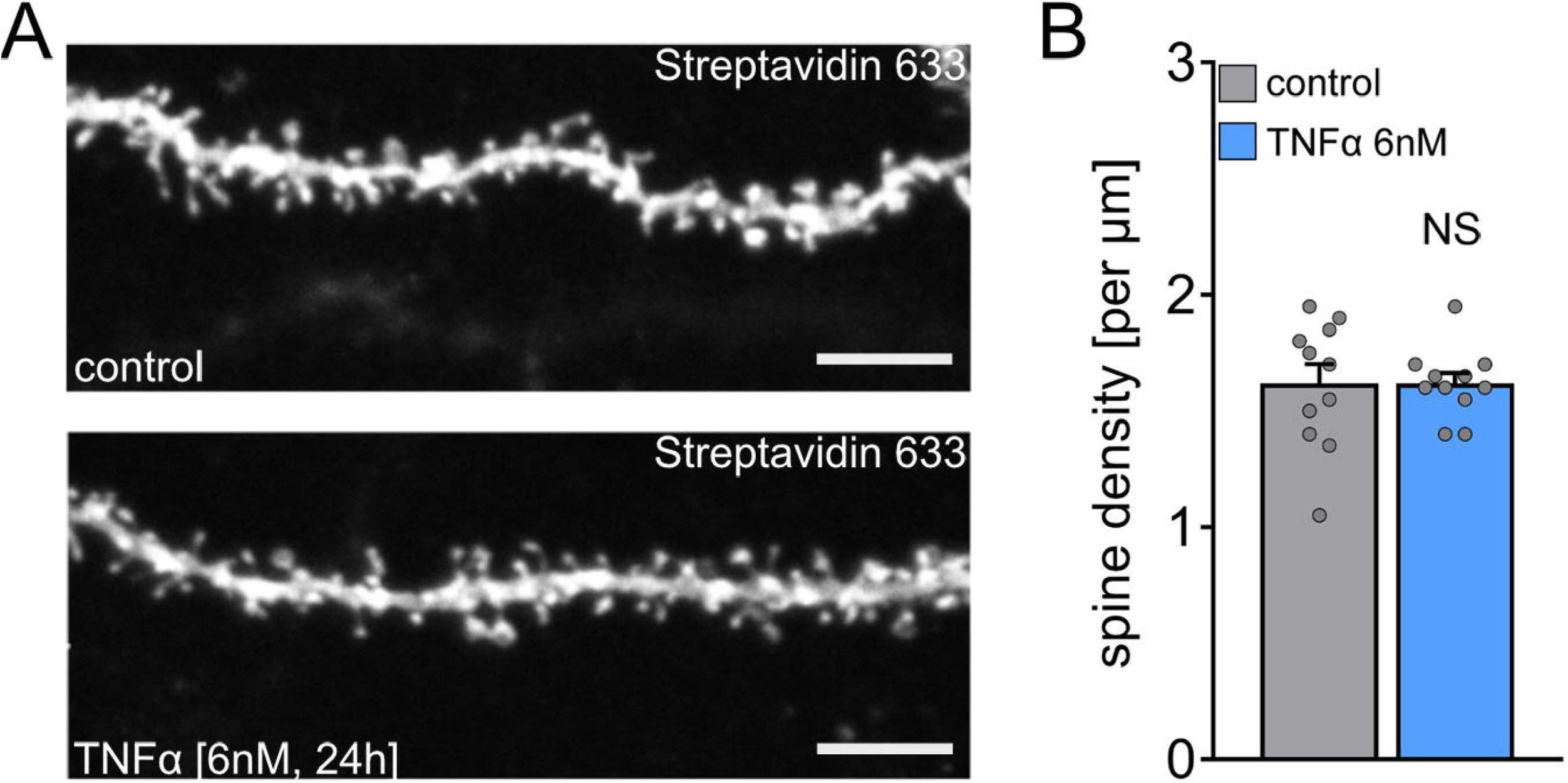
Spine density of CA1 pyramidal neurons is not affected by TNFα. **(A)** Example of dendritic segments of post-hoc-labeled CA1 pyramidal neurons from wild type tissue cultures treated with vehicle-only or TNFα [6 nM, 24]. Scale bar 5 μm. **(B)** Group data for spine densities. (control, n = 11 segments from 11 cells from 4 cultures; TNFα, n = 11 segments from 11 cells from 3 cultures; Mann-Whitney test). Values represent mean ± s.e.m., gray dots indicate individual data points (NS, not significant).

Concordantly, we did not observe any alterations in cell viability in the TNFα group using propidium iodide (PI) staining (Figure 4). PI is a membrane-impermeable agent that does not stain nuclei when membranes are intact [33]. PI signals were comparably low in the control and TNFα-treated tissue cultures, whereas a significant increase in PI signal was observed in the presence of NMDA (50 μM, 24 hours), which served as a positive control. These findings agreed with our results on resting membrane potentials (Figure 1H) and the unaltered spine densities (Figure 3) of CA1 pyramidal neurons in TNFα-treated tissue cultures. We concluded that TNFα (6 nM, 24 hours) increased the GluA1 content of preexisting spine synapses and did not affect cell viability in this setting.

**Figure 4:**
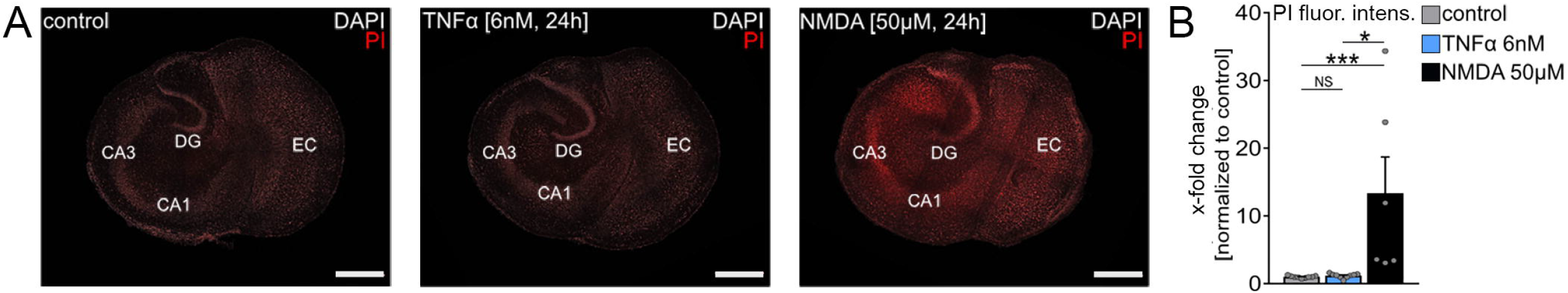
TNFα does not affect cell viability. **(A)** Representative images of tissue cultures treated with vehicle-only or TNFα [6 nM, 24h] and stained with propidium iodide (red) and DAPI (white). NMDA-treated [50 μM, 24h] cultures were used as positive controls. Scale bar 500 μm. **(B)** Group data for propidium iodide stainings. (control, n = 9 cultures; TNFα, n = 9 cultures; NMDA, n = 6 cultures; Kruskal-Wallis test followed by Dunn’s post hoc test). Values represent mean ± s.e.m., gray dots indicate individual data points (*, p<0.05, ***, p<0.001; NS, not significant).

### TNFα induces complex changes in neuronal and microglial gene sets

To further characterize the effects of TNFα, changes in mRNA expression were assessed in vehicle-only and TNFα-treated (6 nM, 24 hours) tissue cultures using transcriptome analysis with Affymetrix chips and qPCR (Figure 5). We found complex changes in several genes for the TNFα group (Figure 5A). Generally applicable gene set enrichment analysis (GSEA, [46]) showed that gene sets related to inflammation were significantly upregulated in TNFα-treated tissue cultures (Figure 5B, shown in red). Specifically, microglial markers, such as *TNFα, IL6, IL1β, CCL2 and CXCL10*, were significantly upregulated (Figure 5C). Interestingly, gene sets associated with various aspects of neuronal physiology, such as neurotransmitter receptors and other synaptic proteins, were significantly downregulated (Figure 5B, shown in blue, 5C).

**Figure 5:**
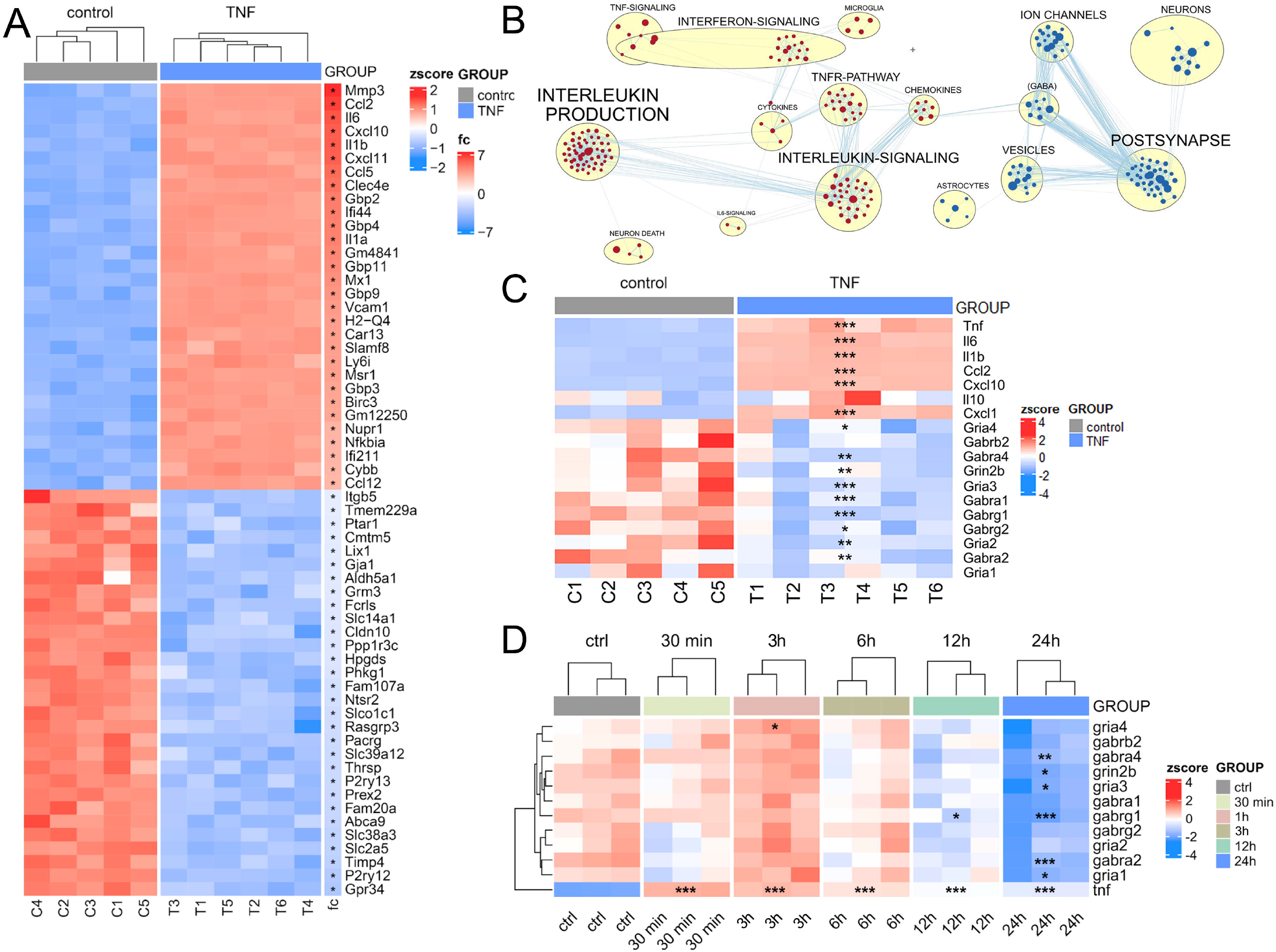
TNFα mediated gene expression changes. **(A)** Heatmap of the 30 most strongly up- and downregulated genes in response to TNFα (6 nM, 24 h). (*, indicates significant difference (adjusted p value < 0.05), linear-model based analysis). **(B)** Enrichment map of gene-sets identified by GSEA; red: up-regulated, blue: downregulated by TNFα treatment. **(C)** Heatmap of gene expression for selected microglial and synaptic markers. **(D)** qPCR analysis of a time-course experiment (after 30 min, 3 h, 6 h, 12 h and 24 h of 6 nM TNFα); heatmap of gene expression for selected genes (*, p<0.05; ***, p<0.01; ***, p<0.001, indicates a significant difference to controls, ANOVA, Tukey’s HSD test corrected for the family-wise error rate).

We confirmed and extended these findings using qPCR analysis (Figure 5D). TNFα-treated tissue cultures were harvested at 30 minutes, 3 hours, 6 hours, 12 hours, and 24 hours. A prominent increase in TNFα mRNA levels was observed at 30 minutes (36.94 ± 1.92-fold vs control), which remained above control levels (5.45 ± 0.18-fold vs control) after 24 hours. Similarly, a trend toward increased mRNA expression of glutamate- and GABA-receptor components was evident within the first 6 hours in the TNFα group, returning to, or below, baseline for some synaptic markers, at 24 hours (Figure 5D). These findings suggest that TNFα rapidly activated microglia and triggered the expression of synaptic proteins. Overall, these data indicated that in the absence of discernible alterations in neural cell viability, as shown with PI staining of TNFα-treated cultures (Figure 4), activated microglia may counter the plasticity-promoting effects of TNFα after a period, consistent with a homeostatic, negative feedback mechanism.

### Microglia are a source of TNFα-induced TNFα production

Next, we tested for the possibility that activated microglia refine TNFα-mediated synaptic plasticity in a homeostatic manner. First, we confirmed that microglia are a major source of TNFα-induced TNFα production (Figure 5C) in tissue cultures prepared from a TNFα-reporter mouse line that expresses green fluorescent protein (GFP) under the control of the TNF promoter (TNF/GFP mice, Figure 6). We recently showed that the GFP signal in tissue cultures prepared from TNF/GFP mice correlates well with changes in TNFα mRNA and TNFα protein levels [33]. Consistent with our microarray analysis, an increase in GFP signal was detected in the microglial cells of TNF/GFP tissue cultures (Figure 6A, B).

**Figure 6:**
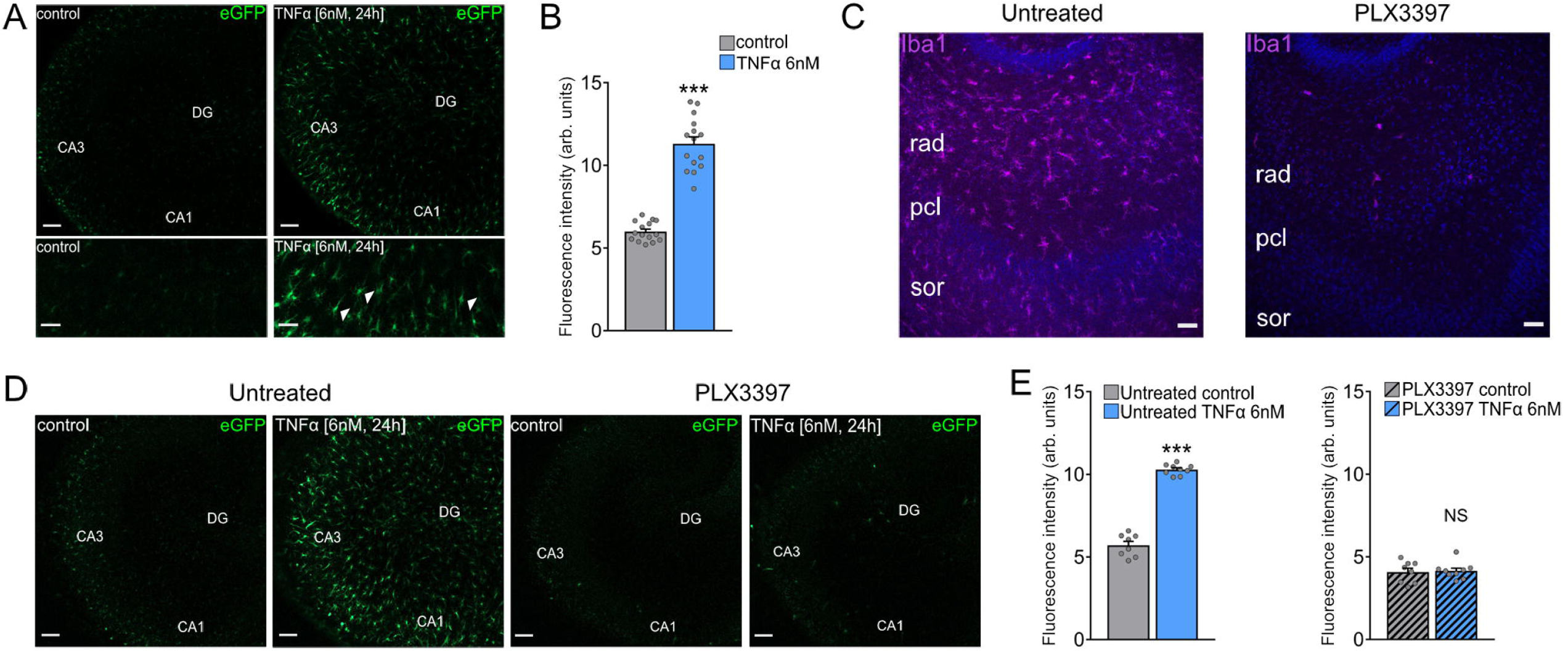
TNFα promotes the production of microglial TNFα. **(A, top)** Representative images from live-cell imaging of tissue cultures prepared from *C57BL/6-Tg(TNFa-eGFP)* mice either treated with vehicle-only (left) or TNFα [6 nM, 24] (right). Scale bar 100 μm. **(A, bottom)** Expanded images of the CA1 region of the same cultures. Microglia shaped cells are indicated with arrowheads. Scale bar 50 μm. **(B)** Group data of GFP fluorescence intensities in vehicle-treated and TNFα [6 nM, 24]-treated cultures. (control, n = 15 cultures; TNFα, n = 15 cultures; Mann-Whitney test). Values represent mean ± s.e.m., gray dots indicate individual data points (***, p<0.001). **(C)** Representative images of tissue cultures treated either with DMSO (upper) or PLX3397 50 nM (lower) and stained against Iba1 to visualize microglia. Scale bar 50 μm. **(D)** Representative images from live-cell imaging of tissue cultures prepared from *C57BL/6-Tg(TNFa-eGFP)* mice either treated with vehicle-only (left) or TNFα [6 nM, 24] (right), after treatment with DMSO or PLX3397 50 nM. Scale bar 100 μm. **(E)** Group data of GFP fluorescence intensities in vehicle-treated and TNFα [6 nM, 24]-treated cultures, after treatment with DMSO or PLX3397 50 nM. (Untreated control, n = 8 cultures; untreated TNFα, n = 9 cultures; PLX3397 control, n = 8 cultures; PLX3397 TNFα, n = 9 cultures; Mann-Whitney test). Values represent mean ± s.e.m., gray dots indicate individual data points. (***, p<0.001; NS, not significant).

We then depleted microglia in tissue cultures prepared from TNF/GFP mice, using the colony stimulating factor 1 receptor (CSF1R) antagonist PLX3397 [31, 47–49]. As shown in Figure 6C, a near-complete depletion of microglia was observed in the presence of 50 nM PLX3397 as indicated by Iba1 staining. In microglia-depleted tissue cultures, there was no significant difference in detected GFP-fluorescence intensity between the TNFα group and vehicle-only controls (Figure 6D, E), confirming that microglia were a major source of TNFα-induced TNFα production.

### TNFα affects excitatory and inhibitory neurotransmission in the absence of microglia

To test for the impact of microglia on TNFα-induced synaptic plasticity, the effects of TNFα (6 nM, 24 hours) on excitatory and inhibitory neurotransmission were probed in microglia-depleted preparations, i.e., in tissue cultures treated with PLX3397 (Figure 7). Consistent with the proposed role of microglia in synaptic homeostasis, a significant increase in the frequency of AMPA receptor-mediated mEPSCs was observed in TNFα-treated cultures, whereas mEPSC amplitude was not significantly different compared to controls (Figure 7A, B). As with tissue cultures containing microglia (Figure 1F), a significant increase in mIPSC frequency of CA1 pyramidal neurons was observed (Figure 7C, D). Therefore, we concluded that TNFα affected both excitatory and inhibitory neurotransmission in the absence of microglia.

**Figure 7:**
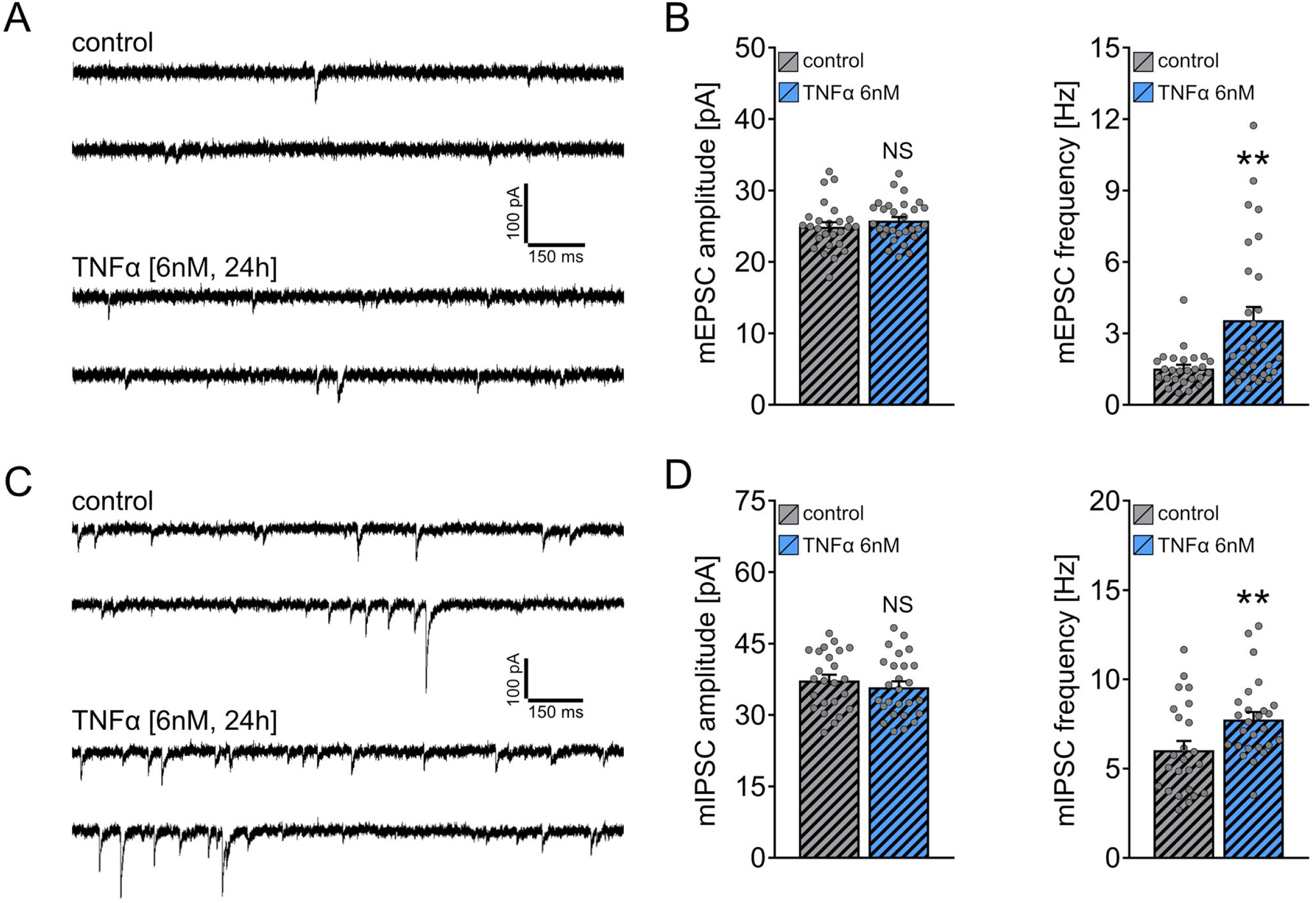
TNFα affects excitatory and inhibitory neurotransmission in the absence of microglia. **(A, B)** Sample traces and group data of AMPA receptor-mediated miniature excitatory postsynaptic currents (mEPSCs) recorded from CA1 pyramidal neurons of microglia-depleted tissue cultures treated with vehicle-only or TNFα [6 nM, 24]. (control, n = 26 cells from 6 cultures; TNFα, n = 29 cells from 6 cultures; Mann-Whitney test). Values represent mean ± s.e.m., gray dots indicate individual data points (**, p<0.01; NS, not significant). **(C, D)** Sample traces and group data of miniature inhibitory postsynaptic currents (mIPSCs) recorded from CA1 pyramidal neurons of microglia-depleted tissue cultures treated with vehicle-only or TNFα [6 nM, 24]. (control, n = 24 cells from 6 cultures; TNFα, n = 26 cells from 6 cultures; Mann-Whitney test). Values represent mean ± s.e.m., gray dots indicate individual data points (**, p<0.01; NS, not significant).

### Low-dose TNFα enhances baseline excitatory neurotransmission without affecting inhibition

After demonstrating that TNFα (6 nM, 24 hours) activated microglia (Figure 5), and increased excitatory synaptic strength in their absence (Figure 7), we reasoned that a lower concentration of TNFα (60 pM), which may not activate microglia, should affect excitatory neurotransmission, increasing mEPSC frequency in microglia-containing wild-type cultures. Notably, we had previously shown that a low concentration (60 pM) of TNFα promoted synaptic plasticity [22].

Wild-type tissue cultures were treated with TNFα (60 pM, 24 hours), and mEPSCs and mIPSCs were recorded from CA1 pyramidal neurons. As shown in Figure 8, a robust increase in mEPSC frequency was observed in the TNFα group (Figure 8A, B). This increase was not observed in the presence of NASPM (10 μM; Figure 8C, D). These results supported our conclusion that TNFα acts by promoting the synaptic accumulation of GluA1. Conversely, mIPSC recordings were indistinguishable between vehicle-only and low TNFα-treated (60 pM, 24 hours) tissue cultures (Figure 8E, F).

**Figure 8:**
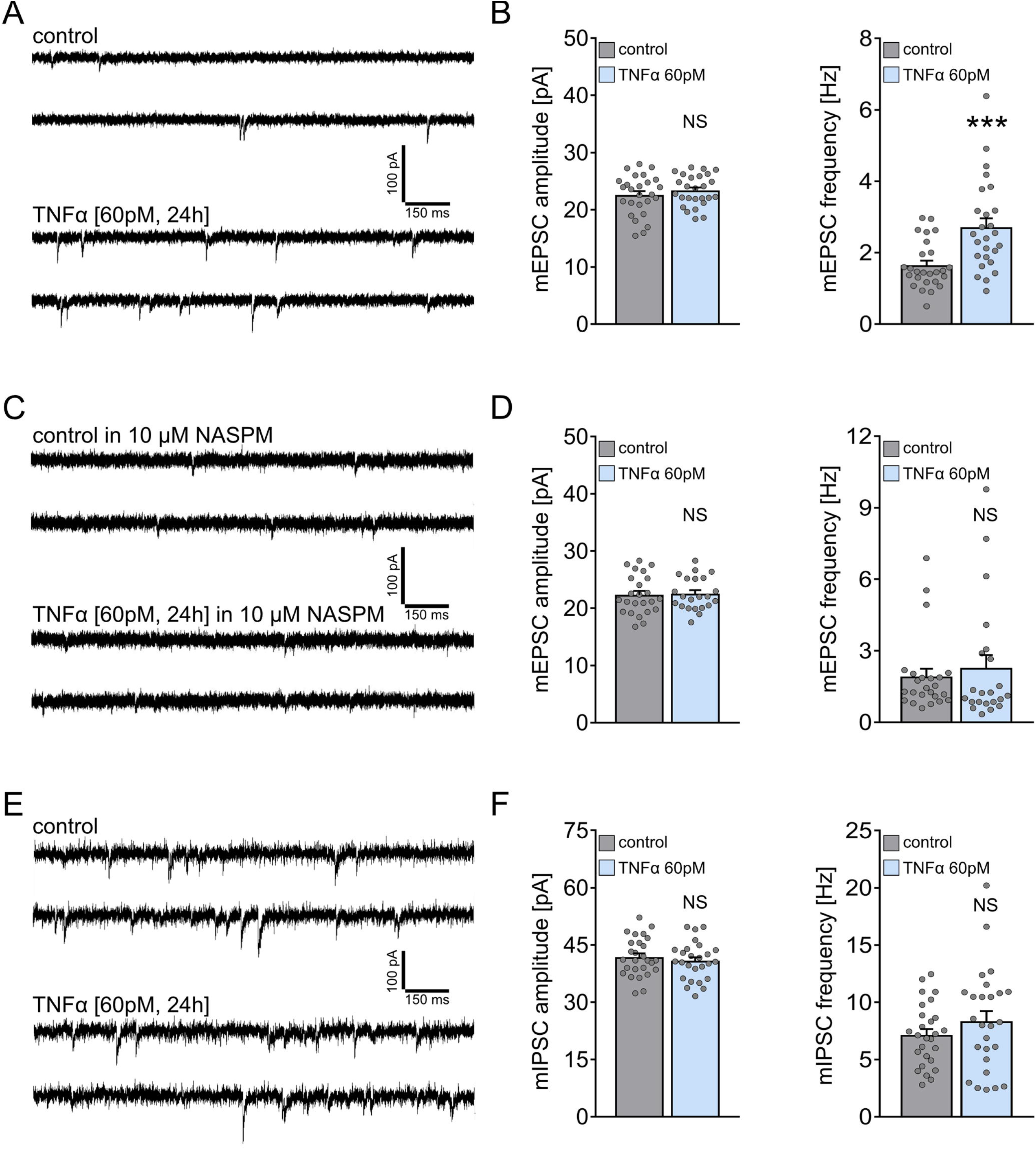
‘Low’ TNFα enhances excitatory neurotransmission without affecting inhibition. **(A, B)** Sample traces and group data of AMPA receptor-mediated miniature excitatory postsynaptic currents (mEPSCs) recorded from CA1 pyramidal neurons of tissue cultures treated with vehicle-only or TNFα [60 pM, 24]. (control, n = 26 cells from 6 cultures; TNFα, n = 26 cells from 6 cultures; Mann-Whitney test). Values represent mean ± s.e.m., gray dots indicate individual data points (***, p<0.001, NS, not significant). **(C, D)** Sample traces and group data of AMPA receptor-mediated miniature excitatory postsynaptic currents (mEPSCs) recorded from CA1 pyramidal neurons of tissue cultures treated with vehicle-only or TNFα [60 pM, 24 in the presence of 1-Naphthylacetyl spermine [NASPM;10 μM]. (control, n = 24 cells from 6 cultures; TNFα, n = 22 cells from 6 cultures; Mann-Whitney test). Values represent mean ± s.e.m., gray dots indicate individual data points (NS, not significant). **(E, F)** Sample traces and group data of miniature inhibitory postsynaptic currents (mIPSCs) recorded from CA1 pyramidal neurons of cultures treated with vehicle-only or TNFα [6 nM, 24]. (control, n = 27 cells from 6 cultures; TNFα, n = 26 cells from 6 cultures; Mann-Whitney test). Values represent mean ± s.e.m., gray dots indicate individual data points (NS, not significant).

Finally, to confirm concentration-dependent effects of TNFα on microglia, we treated another set of tissue cultures with either 60 pM or 6 nM TNFα for 24 hours and determined expression changes for inflammatory cytokines in the incubation media using a protein-detection assay (Figure 9 A; [33]). A significant increase in IL-1β, CXCL1, IL-6, and IL-10 protein levels was only observed in the culture medium of tissue cultures treated with 6 nM, but not those treated with 60 pM, TNFα for 24 hours (Figure 9B-E). These findings suggest that 60 pM TNFα does not activate microglia, while enhancing excitatory, but not inhibitory, neurotransmission. Taken together, we concluded that TNFα induced concentration-dependent effects on inhibitory neurotransmission and that activated microglia, which produced additional TNFα (and other pro- and anti-inflammatory cytokines), are involved in compensating for the plasticity-promoting effects of low TNFα on excitatory postsynapses, i.e., synaptic accumulation of GluA1.

**Figure 9:**
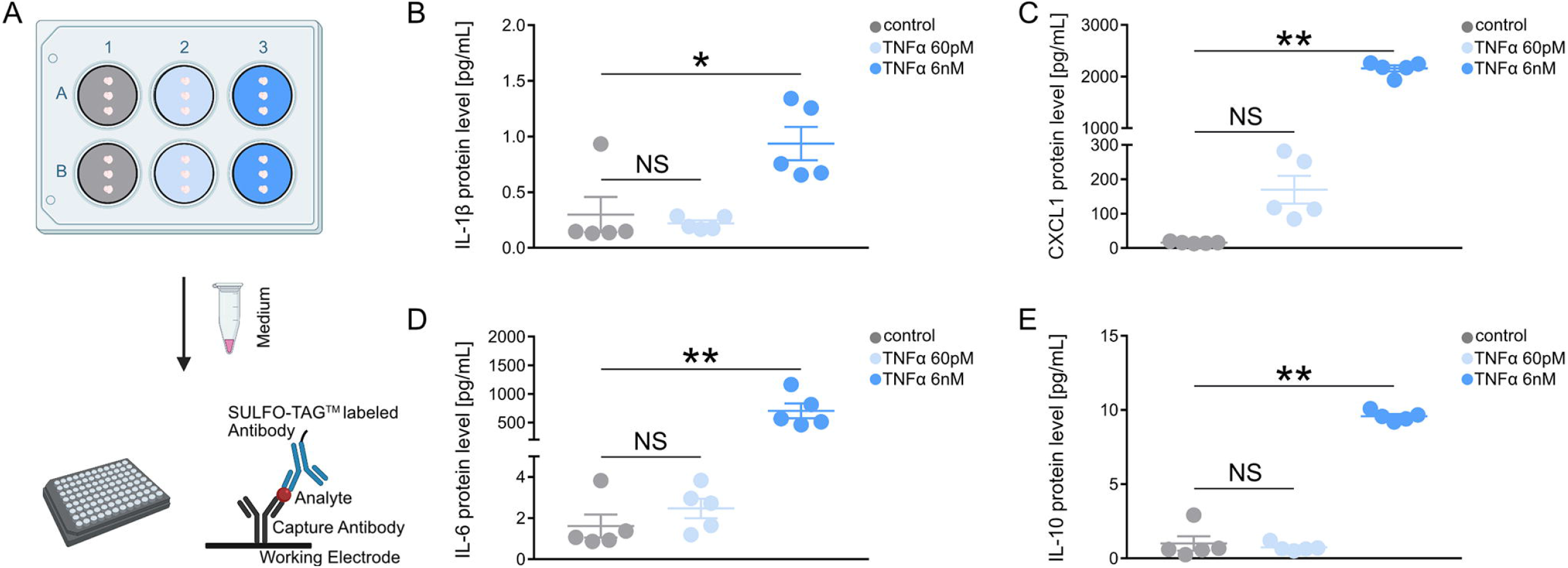
Concentration-dependent effects of TNFα on cytokine release. **(A)** Experimental procedure for determining the expression of cytokines in the culturing medium of vehicle-only or TNFα-treated [60 pM, 24 h or 6 nM, 24h] cultures (3 per filter). Grey: control; Light blue: TNFα [60 pM, 24h]; Blue: TNFα [6 nM, 24h]. 200 μL of culturing medium was collected from each well. A small volume of these samples (analytes) was added to a 96-well plate, pre-coated with capture antibodies against proteins of interest. A solution containing SULFO-TAG™-labeled antibody was added to the wells for detection of the target proteins. **(B-E)** Group data of protein levels in the incubation medium of (B) IL-1β, (C) CXCL1, (D) IL-6 and (E) IL-10 (control, n = 5 samples; TNFα [60 pM, 24h], n = 5 samples; TNFα [6 nM, 24h], n = 5 samples; Kruskal-Wallis test, followed by Dunn’s post hoc test). Values represent mean ± s.e.m., gray dots indicate individual data points. (**, p<0.01; *, p<0.05; NS, not significant).

## DISCUSSION

The results of this study shed new light on the role of TNFα in synaptic plasticity. Consistent with earlier reports we show that TNFα mediates the synaptic accumulation of GluA1– containing AMPA receptors. Interestingly, TNFα modulates synaptic excitation/inhibition balance in a concentration-dependent manner. At low concentrations TNFα increases excitatory neurotransmission; at higher concentrations it increases inhibition. The effects of TNFα on excitatory and inhibitory neurotransmission do not require the presence of microglia since these changes are observed in microglia-depleted tissue cultures as well. In this context, we identified a TNFα-mediated negative feedback mechanism that is based on the activation of microglia, and exerts homeostatic effects on excitatory neurotransmission, thereby emphasizing a role of TNFα-mediated microglia activation in homeostatic synaptic plasticity (Figure 10).

**Figure 10:**
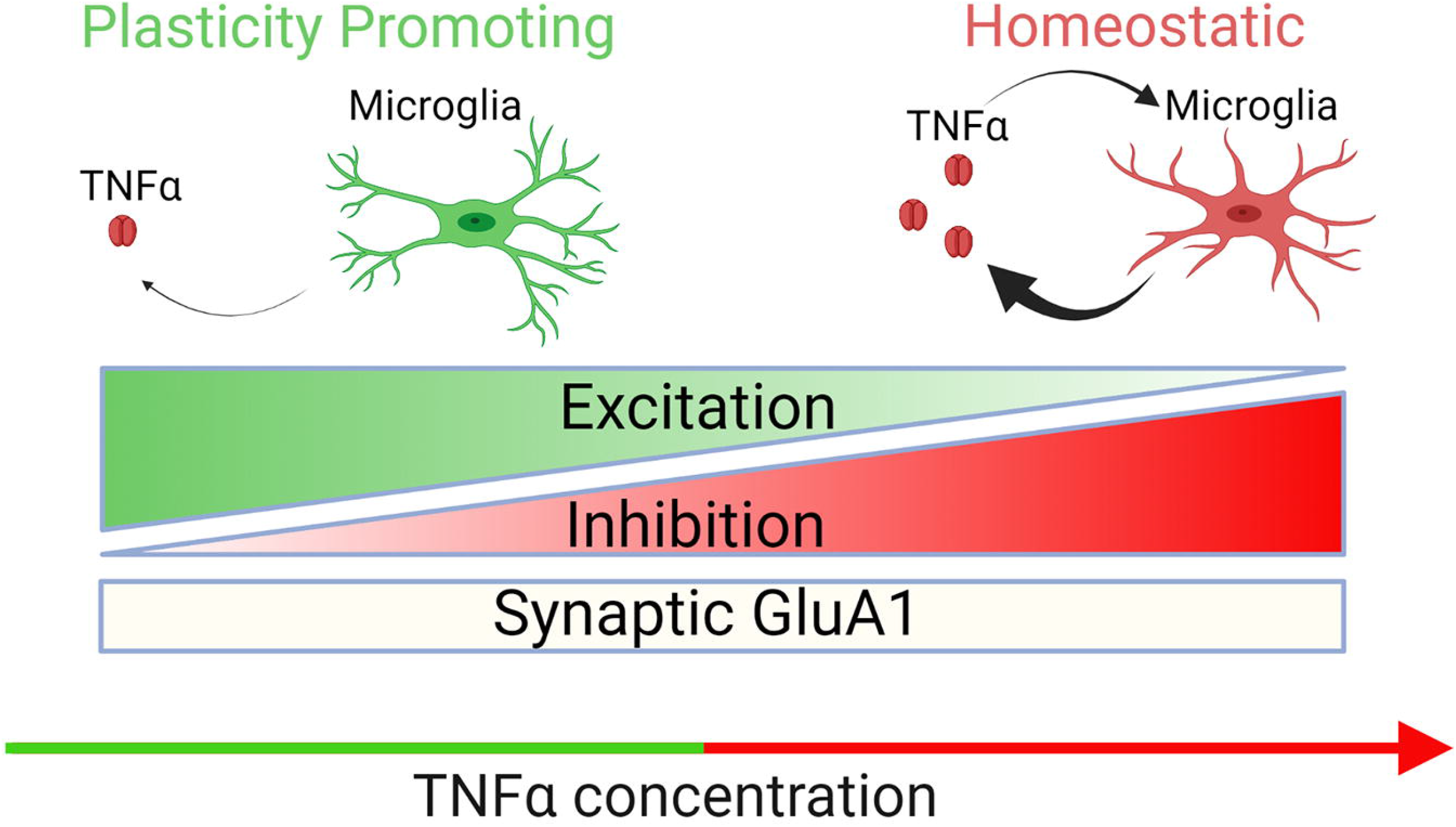
Schematic illustration summarizing the experimental data. TNFα affects synaptic plasticity in a dose-dependent manner, correlated with the activation state of microglia. “Low” TNFα enhances baseline excitatory neurotransmission and increases the number of GluA2-lacking AMPA receptors in the postsynaptic part, while microglia are not activated. Inhibitory neurotransmission is not affected. Increased TNFα concentration activates microglia, which in turn dampens the ability of neurons to express synaptic plasticity, by masking the strengthening of excitatory neurotransmission, most likely in an attempt to prevent excitotoxicity. Nevertheless, the accumulation of GluA2-lacking AMPA receptors takes place normally. Interestingly, microglia are not responsible for the enhanced inhibitory neurotransmission, as inhibitory currents are more frequent after “high” TNFα treatment, both in normal and microglia-depleted tissue cultures.

The first experimental evidence that TNFα affects neurotransmission dates back to 1994, when Grassi et al. showed that short treatment with TNFα increased the frequency of excitatory synaptic currents in dissociated neuronal cultures lacking microglia [50]. This observation attracted considerable interest, particularly after the discovery that TNFα was expressed in the CNS under physiological conditions [7] and the identification of microglia as a major source of TNFα in the brain [9]. It was soon recognized that TNFα affects the ability of neurons to express LTP of excitatory neurotransmission [16, 19, 20]. Our recent studies revealed a complex, concentration-dependent role for TNFα in synaptic plasticity: low concentrations supported the expression of LTP while high concentrations occluded LTP induction [22]. However, it remained unclear how TNFα mediates these concentration-dependent metaplastic effects. Specifically, the role of activated microglia in TNFα-mediated synaptic plasticity had not been previously considered.

The results of this study provide new mechanistic insight on TNFα-mediated synaptic plasticity. At more physiological (low) concentrations, TNFα prompted the synaptic accumulation of GluA1-containing AMPA receptors with no changes in inhibitory neurotransmission observed, consistent with the plasticity promoting effects of TNFα [51–53]. However, TNFα has been also linked to synaptic homeostasis and the adjustment in the balance of synaptic excitation/inhibition [23–26]. Our results suggest that the homeostatic effects of TNFα require the presence of microglia. At higher concentrations, TNFα activated microglia, and no changes in excitatory neurotransmission, as indicated by the unaltered mean mEPSC frequency, were observed (for presynaptic effects of TNFα see [54, 55]). A compensatory reduction in the mRNA levels of some major synaptic markers was evident after 24 hours, while no changes in dendritic spine density were observed. These findings suggest that microglia behave as plasticity-promoting and homeostatic mediators, depending on their activation state. Once TNFα levels exceed a certain concentration, between 60 pM and 6 nM in our experimental setting, activated microglia dampen the plasticity promoting effects of TNFα on excitatory neurotransmission. Notably, we have previously shown that bacterial lipopolysaccharide-induced inflammation, which triggers high levels of microglial TNFα production and other cytokines, blocks the ability of neurons to express synaptic plasticity [33, 56]. The results of the present study suggest that alterations in synaptic plasticity may be regulated by a negative feedback mechanism that prevents the strengthening of excitatory neurotransmission; increased inhibition may contribute to this effect. Whether pathological brain states associated with high TNFα-levels reflect a “hyper”-homeostatic state of disease-associated microglia, which prevent the expression of synaptic plasticity and hamper reorganization and regeneration by mediating “pathological homeostasis”, warrants further investigation [32].

Additional work is also required to clarify the precise mechanisms that trigger the secretion of TNFα under physiological conditions. In this context we were recently able to demonstrate that transcranial magnetic stimulation (TMS), a non-invasive brain stimulation technique used in clinical practice for the treatment of depression and obsessive-compulsive disorders [57, 58], mediates its plasticity promoting effects via microglia activation and cytokine release [49]. It is, therefore, likely that changes in network activity trigger the secretion of plasticity promoting microglial cytokines such as TNFα under physiological conditions.

The mechanisms through which microglial TNFα coordinates the synthesis, synaptic accumulation, and/or degradation of AMPA receptors, both GluA2-containing and GluA2-lacking, and modulates inhibitory neurotransmission in a concentration-dependent manner are not yet fully understood. The effects of TNFα depend upon two distinct receptors, TNFR1 and TNFR2, which are expressed on neurons and glia of the central nervous system [59]. Activation of TNFR1, but not TNFR2, has been linked to the surface accumulation of AMPA receptors [29]. Consistent with this, genetic deletion of TNFR1 (but not TNFR2) led to reduced synaptic levels of AMPA receptors under baseline conditions [42]. Our own work has previously shown that the presence of both TNFR1 and TNFR2 is required for the expression of homeostatic synaptic plasticity [26]. Interestingly, the microarray analysis of the present study showed a significant increase in TNFR2-mRNA, but not TNFR1, in the TNFα group (TNFR1: 1.49 fold change, p = 0.08; TNFR2: 1.95 fold change, p = 0.001). It is, however, difficult to distinguish between neuronal and glial TNFR-signaling pathways and their complex interactions, and the majority of previous studies have used conventional knock-out approaches that remove TNFR1 and/or TNFR2 from all cells of the body (but [55] for astrocytic TNFR1).

Regardless, the results of the present study show that the effects of TNFα on excitatory and inhibitory neurotransmission do not require the presence or activation of microglia, since TNFα-induced changes in mEPSC and mIPSC were observed in microglia-depleted tissue cultures. The homeostatic effects of TNFα on excitatory neurotransmission, however, depend on microglia, with the homeostatic effect and microglial activation not observed in cell cultures exposed to a low concentration. We are confident that future work (1) addressing TNFα-dependent interactions between microglia, astrocytes, and neurons, and (2) identifying the neuronal down-stream mechanisms through which TNFα mediates its concentration-dependent effects on excitatory and inhibitory neurotransmission, will shed further light on the complex role of microglia in health and disease.

## ACKNOWLEDGEMENTS

We would like to thank Gabriele Kaiser, Susanna Glaser and Monika Paetzold for excellent technical support. This work was supported by Deutsche Forschungsgemeinschaft (DFG, Project-ID 259373024 B14 - CRC/TRR 167). MB is supported by SFB1479–Project-ID 441891347-S1, SFB1453–Project-ID 431984000-S1 and CRC/TRR167–Project-ID Z01). MB and GA are supported by the German Federal Ministry of Education and Research by MIRACUM within the Medical Informatics Funding Scheme (FKZ 01ZZ1801B for MB and EkoEstMed-FKZ 01ZZ2015 for GA).

## REFERENCES

1. Becher, B., S. Spath, and J. Goverman. Cytokine networks in neuroinflammation. Nat Rev Immunol, 2017, 17, 49–59. DOI: 10.1038/nri.2016.123.

2. Yang, Q. Q., and J. W. Zhou. Neuroinflammation in the central nervous system: Symphony of glial cells. Glia, 2019, 67, 1017–1035. DOI: 10.1002/glia.23571.

3. Glass, C. K., K. Saijo, B. Winner, M. C. Marchetto, and F. H. Gage. Mechanisms underlying inflammation in neurodegeneration. Cell, 2010, 140, 918–934. DOI: 10.1016/j.cell.2010.02.016.

4. Kempuraj, D., R. Thangavel, P. A. Natteru, G. P. Selvakumar, D. Saeed, H. Zahoor, S. Zaheer, S. S. Iyer, and A. Zaheer. Neuroinflammation Induces Neurodegeneration. J Neurol Neurosurg Spine, 2016, 1. DOI.

5. Stephenson, J., E. Nutma, P. van der Valk, and S. Amor. Inflammation in CNS neurodegenerative diseases. Immunology, 2018, 154, 204–219. DOI: 10.1111/imm.12922.

6. Kwon, H. S., and S. H. Koh. Neuroinflammation in neurodegenerative disorders: the roles of microglia and astrocytes. Transl Neurodegener, 2020, 9, 42. DOI: 10.1186/s40035-020-00221-2.

7. Vitkovic, L., J. Bockaert, and C. Jacque. “Inflammatory” cytokines: neuromodulators in normal brain? J Neurochem, 2000, 74, 457–471. DOI: 10.1046/j.1471-4159.2000.740457.x.

8. Bourgognon, J. M., and J. Cavanagh. The role of cytokines in modulating learning and memory and brain plasticity. Brain Neurosci Adv, 2020, 4, 2398212820979802. DOI: 10.1177/2398212820979802.

9. Hanisch, U. K. Microglia as a source and target of cytokines. Glia, 2002, 40, 140–155. DOI: 10.1002/glia.10161.

10. Goldmann, T., P. Wieghofer, P. F. Muller, Y. Wolf, D. Varol, S. Yona, S. M. Brendecke, K. Kierdorf, O. Staszewski, M. Datta, T. Luedde, M. Heikenwalder, S. Jung, and M. Prinz. A new type of microglia gene targeting shows TAK1 to be pivotal in CNS autoimmune inflammation. Nat Neurosci, 2013, 16, 1618–1626. DOI: 10.1038/nn.3531.

11. Welser-Alves, J. V., and R. Milner. Microglia are the major source of TNF-alpha and TGF-beta1 in postnatal glial cultures; regulation by cytokines, lipopolysaccharide, and vitronectin. Neurochem Int, 2013, 63, 47–53. DOI: 10.1016/j.neuint.2013.04.007.

12. Prinz, M., T. Masuda, M. A. Wheeler, and F. J. Quintana. Microglia and Central Nervous System-Associated Macrophages-From Origin to Disease Modulation. Annu Rev Immunol, 2021, 39, 251–277. DOI: 10.1146/annurev-immunol-093019-110159.

13. Yirmiya, R., and I. Goshen. Immune modulation of learning, memory, neural plasticity and neurogenesis. Brain Behav Immun, 2011, 25, 181–213. DOI: 10.1016/j.bbi.2010.10.015.

14. Levin, S. G., and O. V. Godukhin. Modulating Effect of Cytokines on Mechanisms of Synaptic Plasticity in the Brain. Biochemistry (Mosc), 2017, 82, 264–274. DOI: 10.1134/S000629791703004X.

15. Nistico, R., E. Salter, C. Nicolas, M. Feligioni, D. Mango, Z. A. Bortolotto, P. Gressens, G. L. Collingridge, and S. Peineau. Synaptoimmunology - roles in health and disease. Mol Brain, 2017, 10, 26. DOI: 10.1186/s13041-017-0308-9.

16. Cunningham, A. J., C. A. Murray, L. A. O’Neill, M. A. Lynch, and J. J. O’Connor. Interleukin-1 beta (IL-1 beta) and tumour necrosis factor (TNF) inhibit long-term potentiation in the rat dentate gyrus in vitro. Neurosci Lett, 1996, 203, 17–20. DOI: 10.1016/0304-3940(95)12252-4.

17. Albensi, B. C., and M. P. Mattson. Evidence for the involvement of TNF and NF-kappaB in hippocampal synaptic plasticity. Synapse, 2000, 35, 151–159. DOI: 10.1002/(SICI)1098-2396(200002)35:2<151::AID-SYN8>3.0.CO;2-P.

18. Curran, B. P., and J. J. O’Connor. The inhibition of long-term potentiation in the rat dentate gyrus by pro-inflammatory cytokines is attenuated in the presence of nicotine. Neurosci Lett, 2003, 344, 103–106. DOI: 10.1016/s0304-3940(03)00440-3.

19. Butler, M. P., J. J. O’Connor, and P. N. Moynagh. Dissection of tumor-necrosis factor-alpha inhibition of long-term potentiation (LTP) reveals a p38 mitogen-activated protein kinase-dependent mechanism which maps to early-but not late-phase LTP. Neuroscience, 2004, 124, 319–326. DOI: 10.1016/j.neuroscience.2003.11.040.

20. Wall, A. M., G. Mukandala, N. H. Greig, and J. J. O’Connor. Tumor necrosis factor-alpha potentiates long-term potentiation in the rat dentate gyrus after acute hypoxia. J Neurosci Res, 2015, 93, 815–829. DOI: 10.1002/jnr.23540.

21. Rizzo, F. R., A. Musella, F. De Vito, D. Fresegna, S. Bullitta, V. Vanni, L. Guadalupi, M. Stampanoni Bassi, F. Buttari, G. Mandolesi, D. Centonze, and A. Gentile. Tumor Necrosis Factor and Interleukin-1beta Modulate Synaptic Plasticity during Neuroinflammation. Neural Plast, 2018, 2018, 8430123. DOI: 10.1155/2018/8430123.

22. Maggio, N., and A. Vlachos. Tumor necrosis factor (TNF) modulates synaptic plasticity in a concentration-dependent manner through intracellular calcium stores. J Mol Med (Berl), 2018, 96, 1039–1047. DOI: 10.1007/s00109-018-1674-1.

23. Stellwagen, D., and R. C. Malenka. Synaptic scaling mediated by glial TNF-alpha. Nature, 2006, 440, 1054–1059. DOI: 10.1038/nature04671.

24. Steinmetz, C. C., and G. G. Turrigiano. Tumor necrosis factor-alpha signaling maintains the ability of cortical synapses to express synaptic scaling. J Neurosci, 2010, 30, 14685–14690. DOI: 10.1523/JNEUROSCI.2210-10.2010.

25. Becker, D., N. Zahn, T. Deller, and A. Vlachos. Tumor necrosis factor alpha maintains denervation-induced homeostatic synaptic plasticity of mouse dentate granule cells. Front Cell Neurosci, 2013, 7, 257. DOI: 10.3389/fncel.2013.00257.

26. Becker, D., T. Deller, and A. Vlachos. Tumor necrosis factor (TNF)-receptor 1 and 2 mediate homeostatic synaptic plasticity of denervated mouse dentate granule cells. Sci Rep, 2015, 5, 12726. DOI: 10.1038/srep12726.

27. Singh, A., O. D. Jones, B. G. Mockett, S. M. Ohline, and W. C. Abraham. Tumor Necrosis Factor-alpha-Mediated Metaplastic Inhibition of LTP Is Constitutively Engaged in an Alzheimer’s Disease Model. J Neurosci, 2019, 39, 9083–9097. DOI: 10.1523/JNEUROSCI.1492-19.2019.

28. Singh, A., S. Sateesh, O. D. Jones, and W. C. Abraham. Pathway-specific TNF-mediated metaplasticity in hippocampal area CA1. Sci Rep, 2022, 12, 1746. DOI: 10.1038/s41598-022-05844-1.

29. Stellwagen, D., E. C. Beattie, J. Y. Seo, and R. C. Malenka. Differential regulation of AMPA receptor and GABA receptor trafficking by tumor necrosis factor-alpha. J Neurosci, 2005, 25, 3219–3228. DOI: 10.1523/JNEUROSCI.4486-04.2005.

30. Pribiag, H., and D. Stellwagen. TNF-alpha downregulates inhibitory neurotransmission through protein phosphatase 1-dependent trafficking of GABA(A) receptors. J Neurosci, 2013, 33, 15879–15893. DOI: 10.1523/JNEUROSCI.0530-13.2013.

31. Kleidonas, D., and A. Vlachos. Scavenging Tumor Necrosis Factor alpha Does Not Affect Inhibition of Dentate Granule Cells Following In Vitro Entorhinal Cortex Lesion. Cells, 2021, 10. DOI: 10.3390/cells10113232.

32. Galanis, C., and A. Vlachos. Hebbian and Homeostatic Synaptic Plasticity-Do Alterations of One Reflect Enhancement of the Other? Front Cell Neurosci, 2020, 14, 50. DOI: 10.3389/fncel.2020.00050.

33. Lenz, M., A. Eichler, P. Kruse, A. Strehl, S. Rodriguez-Rozada, I. Goren, N. Yogev, S. Frank, A. Waisman, T. Deller, S. Jung, N. Maggio, and A. Vlachos. Interleukin 10 Restores Lipopolysaccharide-Induced Alterations in Synaptic Plasticity Probed by Repetitive Magnetic Stimulation. Front Immunol, 2020, 11, 614509. DOI: 10.3389/fimmu.2020.614509.

34. Del Turco, D., and T. Deller. Organotypic entorhino-hippocampal slice cultures--a tool to study the molecular and cellular regulation of axonal regeneration and collateral sprouting in vitro. Methods Mol Biol, 2007, 399, 55–66. DOI: 10.1007/978-1-59745-504-6_5.

35. Ritchie, M. E., B. Phipson, D. Wu, Y. Hu, C. W. Law, W. Shi, and G. K. Smyth. limma powers differential expression analyses for RNA-sequencing and microarray studies. Nucleic Acids Res, 2015, 43, e47. DOI: 10.1093/nar/gkv007.

36. Mootha, V. K., C. M. Lindgren, K. F. Eriksson, A. Subramanian, S. Sihag, J. Lehar, P. Puigserver, E. Carlsson, M. Ridderstrale, E. Laurila, N. Houstis, M. J. Daly, N. Patterson, J. P. Mesirov, T. R. Golub, P. Tamayo, B. Spiegelman, E. S. Lander, J. N. Hirschhorn, D. Altshuler, and L. C. Groop. PGC-1alpha-responsive genes involved in oxidative phosphorylation are coordinately downregulated in human diabetes. Nat Genet, 2003, 34, 267–273. DOI: 10.1038/ng1180.

37. Subramanian, A., P. Tamayo, V. K. Mootha, S. Mukherjee, B. L. Ebert, M. A. Gillette, A. Paulovich, S. L. Pomeroy, T. R. Golub, E. S. Lander, and J. P. Mesirov. Gene set enrichment analysis: a knowledge-based approach for interpreting genome-wide expression profiles. Proc Natl Acad Sci U S A, 2005, 102, 15545–15550. DOI: 10.1073/pnas.0506580102.

38. Shannon, P., A. Markiel, O. Ozier, N. S. Baliga, J. T. Wang, D. Ramage, N. Amin, B. Schwikowski, and T. Ideker. Cytoscape: a software environment for integrated models of biomolecular interaction networks. Genome Res, 2003, 13, 2498–2504. DOI: 10.1101/gr.1239303.

39. Vlachos, A., F. Muller-Dahlhaus, J. Rosskopp, M. Lenz, U. Ziemann, and T. Deller. Repetitive magnetic stimulation induces functional and structural plasticity of excitatory postsynapses in mouse organotypic hippocampal slice cultures. J Neurosci, 2012, 32, 17514–17523. DOI: 10.1523/JNEUROSCI.0409-12.2012.

40. Beattie, E. C., D. Stellwagen, W. Morishita, J. C. Bresnahan, B. K. Ha, M. Von Zastrow, M. S. Beattie, and R. C. Malenka. Control of synaptic strength by glial TNFalpha. Science, 2002, 295, 2282–2285. DOI: 10.1126/science.1067859.

41. Ogoshi, F., H. Z. Yin, Y. Kuppumbatti, B. Song, S. Amindari, and J. H. Weiss. Tumor necrosis-factor-alpha (TNF-alpha) induces rapid insertion of Ca2+-permeable alpha-amino-3-hydroxyl-5-methyl-4-isoxazole-propionate (AMPA)/kainate (Ca-A/K) channels in a subset of hippocampal pyramidal neurons. Exp Neurol, 2005, 193, 384–393. DOI: 10.1016/j.expneurol.2004.12.026.

42. He, P., Q. Liu, J. Wu, and Y. Shen. Genetic deletion of TNF receptor suppresses excitatory synaptic transmission via reducing AMPA receptor synaptic localization in cortical neurons. FASEB J, 2012, 26, 334–345. DOI: 10.1096/fj.11-192716.

43. Carvalho, T. P., and D. V. Buonomano. Differential effects of excitatory and inhibitory plasticity on synaptically driven neuronal input-output functions. Neuron, 2009, 61, 774–785. DOI: 10.1016/j.neuron.2009.01.013.

44. Wang, L., and A. Maffei. Inhibitory plasticity dictates the sign of plasticity at excitatory synapses. J Neurosci, 2014, 34, 1083–1093. DOI: 10.1523/JNEUROSCI.4711-13.2014.

45. Man, H. Y. GluA2-lacking, calcium-permeable AMPA receptors--inducers of plasticity? Curr Opin Neurobiol, 2011, 21, 291–298. DOI: 10.1016/j.conb.2011.01.001.

46. Luo, W., M. S. Friedman, K. Shedden, K. D. Hankenson, and P. J. Woolf. GAGE: generally applicable gene set enrichment for pathway analysis. BMC Bioinformatics, 2009, 10, 161. DOI: 10.1186/1471-2105-10-161.

47. Han, J., R. A. Harris, and X. M. Zhang. An updated assessment of microglia depletion: current concepts and future directions. Mol Brain, 2017, 10, 25. DOI: 10.1186/s13041-017-0307-x.

48. Coleman, L. G., Jr., J. Zou, and F. T. Crews. Microglial depletion and repopulation in brain slice culture normalizes sensitized proinflammatory signaling. J Neuroinflammation, 2020, 17, 27. DOI: 10.1186/s12974-019-1678-y.

49. Eichler, Amelie, Dimitrios Kleidonas, Zsolt Turi, Matthias Kirsch, Dietmar Pfeifer, Takahiro Masuda, Marco Prinz, Maximilian Lenz, and Andreas Vlachos. Microglia mediate synaptic plasticity induced by 10 Hz repetitive magnetic stimulation. bioRxiv, 2021, 2021.2010.2003.462905. DOI: 10.1101/2021.10.03.462905.

50. Grassi, F., A. M. Mileo, L. Monaco, A. Punturieri, A. Santoni, and F. Eusebi. TNF-alpha increases the frequency of spontaneous miniature synaptic currents in cultured rat hippocampal neurons. Brain Res, 1994, 659, 226–230. DOI: 10.1016/0006-8993(94)90883-4.

51. Hu, H., E. Real, K. Takamiya, M. G. Kang, J. Ledoux, R. L. Huganir, and R. Malinow. Emotion enhances learning via norepinephrine regulation of AMPA-receptor trafficking. Cell, 2007, 131, 160–173. DOI: 10.1016/j.cell.2007.09.017.

52. Tenorio, G., S. A. Connor, D. Guevremont, W. C. Abraham, J. Williams, T. J. O’Dell, and P. V. Nguyen. ‘Silent’ priming of translation-dependent LTP by ss-adrenergic receptors involves phosphorylation and recruitment of AMPA receptors. Learn Mem, 2010, 17, 627–638. DOI: 10.1101/lm.1974510.

53. Whitehead, G., J. Jo, E. L. Hogg, T. Piers, D. H. Kim, G. Seaton, H. Seok, G. Bru-Mercier, G. H. Son, P. Regan, L. Hildebrandt, E. Waite, B. C. Kim, T. L. Kerrigan, K. Kim, D. J. Whitcomb, G. L. Collingridge, S. L. Lightman, and K. Cho. Acute stress causes rapid synaptic insertion of Ca2+ -permeable AMPA receptors to facilitate long-term potentiation in the hippocampus. Brain, 2013, 136, 3753–3765. DOI: 10.1093/brain/awt293.

54. Santello, M., P. Bezzi, and A. Volterra. TNFalpha controls glutamatergic gliotransmission in the hippocampal dentate gyrus. Neuron, 2011, 69, 988–1001. DOI: 10.1016/j.neuron.2011.02.003.

55. Habbas, S., M. Santello, D. Becker, H. Stubbe, G. Zappia, N. Liaudet, F. R. Klaus, G. Kollias, A. Fontana, C. R. Pryce, T. Suter, and A. Volterra. Neuroinflammatory TNFalpha Impairs Memory via Astrocyte Signaling. Cell, 2015, 163, 1730–1741. DOI: 10.1016/j.cell.2015.11.023.

56. Strehl, A., M. Lenz, Z. Itsekson-Hayosh, D. Becker, J. Chapman, T. Deller, N. Maggio, and A. Vlachos. Systemic inflammation is associated with a reduction in Synaptopodin expression in the mouse hippocampus. Exp Neurol, 2014, 261, 230–235. DOI: 10.1016/j.expneurol.2014.04.033.

57. George, M. S., and R. M. Post. Daily left prefrontal repetitive transcranial magnetic stimulation for acute treatment of medication-resistant depression. Am J Psychiatry, 2011, 168, 356–364. DOI: 10.1176/appi.ajp.2010.10060864.

58. Cocchi, L., A. Zalesky, Z. Nott, G. Whybird, P. B. Fitzgerald, and M. Breakspear. Transcranial magnetic stimulation in obsessive-compulsive disorder: A focus on network mechanisms and state dependence. Neuroimage Clin, 2018, 19, 661–674. DOI: 10.1016/j.nicl.2018.05.029.

59. Santello, M., and A. Volterra. TNFalpha in synaptic function: switching gears. Trends Neurosci, 2012, 35, 638–647. DOI: 10.1016/j.tins.2012.06.001.

